# Stem cell mitotic drive ensures asymmetric epigenetic inheritance

**DOI:** 10.1101/416446

**Authors:** Rajesh Ranjan, Jonathan Snedeker, Xin Chen

**Author notes:** Correspondence to: Xin Chen, Ph.D., Department of Biology, 3400 North Charles Street, The Johns Hopkins University, Baltimore, MD 21218-2685.

## Abstract

During *Drosophila* male germline stem cell asymmetric division, sister centromeres communicate with spindle microtubules for differential attachment and segregation of sister chromatids.

**SUMMARY:** Through the process of symmetric cell division, one mother cell gives rise to two identical daughter cells. Many stem cells utilize asymmetric cell division (ACD) to produce a self-renewed stem cell and a differentiating daughter cell. Since both daughter cells inherit the identical genetic information during ACD, a crucial question concerns how non-genic factors could be inherited differentially to establish distinct cell fates. It has been hypothesized that epigenetic differences at sister centromeres could contribute to biased sister chromatid attachment and segregation. However, direct *in vivo* evidence has never been shown. Here, we report that a stem cell-specific ‘mitotic drive’ ensures biased sister chromatid attachment and segregation. We have found during stem cell ACD, sister centromeres become asymmetrically enriched with proteins involved in centromere specification and kinetochore function. Furthermore, we show that that temporally asymmetric microtubule activities direct polarized nuclear envelope breakdown, allowing for the preferential recognition and attachment of microtubules to asymmetric sister kinetochores and sister centromeres. This communication occurs in a spatiotemporally regulated manner. Abolishment of either the establishment of asymmetric sister centromeres or the asymmetric microtubule emanation results in randomized sister chromatid segregation, which leads to stem cell loss. Our results demonstrate that the *cis*-asymmetry at sister centromeres tightly coordinates with the *trans*-asymmetry from the mitotic machinery to allow for differential attachment and segregation of genetically identical yet epigenetically distinct sister chromatids. Together, these results provide the first direct *in vivo* mechanisms for partitioning epigenetically distinct sister chromatids in asymmetrically dividing stem cells, which opens a new direction to study how this mechanism could be used in other developmental contexts to achieve distinct cell fates through mitosis.

## INTRODUCTION

Epigenetic mechanisms play important roles in regulating cell identity and function. Mis-regulation of epigenetic information may lead to abnormalities in cellular behaviors that underlie many diseases, such as developmental defects, cancer, and tissue degeneration (Allis and Jenuwein, 2016). Histone proteins, which play important roles in DNA packaging and chromosomal structure, represent a major group of epigenetic information carriers (Sitbon et al., 2017).

Many types of adult stem cells undergo asymmetric cell division (ACD) to generate both a self-renewed stem cell and a daughter cell, which will subsequently differentiate (Clevers, 2005; Kahney et al., 2017; Knoblich, 2010; Morrison and Kimble, 2006; Venkei and Yamashita, 2018). Since stem cells have unique gene expression [e.g. (Amcheslavsky et al., 2014; Blanpain and Fuchs, 2006; Kai et al., 2005; Seale et al., 2004; Terry et al., 2006; Terskikh et al., 2003; Young, 2011)], the field has long sought to understand how stem cells could maintain their epigenetic information and gene expression through many cell divisions.

During the asymmetric division of *Drosophila* male germline stem cells (GSCs) (Figure 1A), we showed previously that the preexisting (old) histone H3 is selectively segregated to the GSC, whereas the newly synthesized (new) H3 is enriched in the differentiating daughter cell known as a gonialblast (GB) (Tran et al., 2012). This asymmetry is unique to stem cells, as symmetrically dividing spermatogonial cells (SGs) show symmetric patterns of histone H3 inheritance. We also identified that differential phosphorylation at Threonine 3 of histone H3 (H3T3P) distinguishes old *versus* new H3 in the asymmetrically dividing GSCs. Mis-regulation of this phosphorylation leads to randomized segregation of old H3 *versus* new H3, as well as stem cell loss and early-stage germline tumors (Xie et al., 2015). We hypothesize that prior to mitosis, old H3 and new H3 are differentially distributed at the two sets of sister chromatids (Wooten, 2018). During the subsequent mitosis, the two sets of epigenetically distinct sister chromatids are asymmetrically segregated (Tran et al., 2013; Xie et al., 2017). According to this model, sister chromatids carrying distinct epigenetic information need to communicate with the mitotic machinery to achieve differential attachment, followed by asymmetric segregation. It has been shown previously that centrosomes display asymmetry in *Drosophila* male GSCs, wherein the mother centrosome is retained by the GSC while the daughter centrosome is inherited by the GB (Yamashita et al., 2007). However, it remains unclear whether this asymmetry in centrosome inheritance is related to the phenomenon of asymmetric histone inheritance.

**Figure 1:**
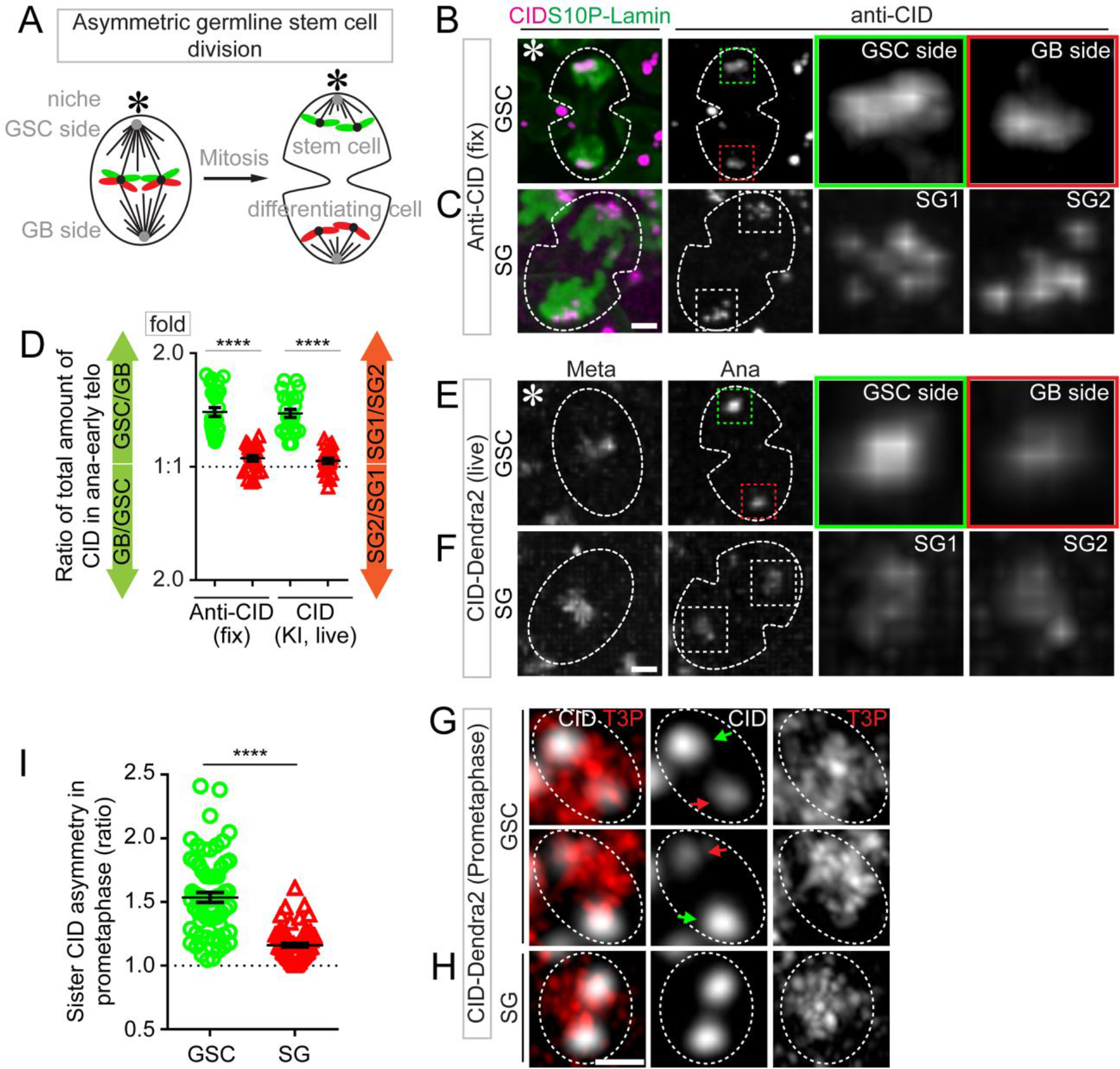
Asymmetric CID inheritance in *Drosophila* male GSCs. (**A**) A cartoon depicting asymmetric *Drosophila* male GSC division and asymmetric histone H3 inheritance (old H3 - green, new H3 - red). (B-F) Asymmetric CID segregation in asymmetrically dividing GSCs with more CID toward the GSC side, using both immunostaining of fixed cells (B) and live cell imaging of a CID-Dendra2 knock-in line (E). In symmetric SG cell division, CID is symmetrically distributed, as determined using both immunostaining (C) and live cell imaging (F). Quantification of all four data sets in (D): 1.41± 0.04-fold for GSC/GB (*n* = 27), 1.06± 0.02-fold for SG1/SG2 (*n* = 23), as determined by anti-CID, Table S1; 1.39± 0.04-fold for GSC/GB (*n* = 23), 1.04± 0.02-fold for SG1/SG2 (*n* = 21), as determined by live cell imaging using CID-Dendra2, Table S2. (G-H) In prometaphase, resolved individual sister centromeres show more asymmetry in GSCs [green arrow-stronger centromere, red arrow-weak centromere, (G)], compared with resolved sister centromeres in SGs (H). (I) Quantification of individual pairs of resolved sister centromeres in GSCs (1.54± 0.04-fold, *n* = 65) and SGs (1.16± 0.02-fold, *n* = 66) at prometaphase, Table S4. All ratios = Avg± SE. *P*-value: paired *t* test. ****: *P* < 10^−4^. Asterisk: hub. Scale bars: 2μm.

During mitosis, microtubules emanate from centrosomes and attach to sister chromatids at the centromeric region through the kinetochore protein complex (Cheeseman, 2014; Fukagawa and Earnshaw, 2014; Rieder, 2005). Centromeres are specialized regions of chromosomes which are often defined by epigenetic characteristics (Bodor et al., 2014; Dunleavy et al., 2005; Karpen and Allshire, 1997; Schueler and Sullivan, 2006; Vos et al., 2006; Westhorpe and Straight, 2014). In most eukaryotic cells, the critical epigenetic feature of centromeres is the centromere-specific histone H3 variant, known as the Centromere identifier (CID) in flies (Henikoff et al., 2000) and Centromere protein A (CENP-A) in mammals (Allshire and Karpen, 2008; Mendiburo et al., 2011; Palmer et al., 1987). Both are structurally different from the canonical H3 (McKinley and Cheeseman, 2016; Melters et al., 2015; Miell et al., 2013). Based on the unique features and essential roles of centromeres in coordinating sister chromatid segregation during mitosis, it has been proposed that epigenetic differences between sister centromeres could ensure that stem cells retain their unique epigenetic information and gene expression, even after many cell divisions (Lansdorp, 2007). However, direct *in vivo* evidence has never been shown.

## RESULTS

### Asymmetric Sister Centromere Inheritance in Male GSCs

During *Drosophila* male GSC asymmetric division at anaphase or early telophase, the level of endogenous CID showed a 1.41-fold enrichment at sister chromatids segregating toward the future GSC side compared to the future GB side, stained by anti-CID antibodies (Figures 1B and 1D, Figures S1A and S1C). Conversely, symmetrically dividing SGs showed nearly equal CID distribution at anaphase or early telophase (Figures 1C and 1D, Figures S1B and S1D). To confirm this result, live cell imaging was performed using a knock-in fly strain with the endogenous *cid* gene tagged with a fluorescent protein-encoding *Dendra2* (Chudakov et al., 2007), which was generated by CRISPR/Cas9-mediated genome editing (Horvath and Barrangou, 2010; Wright et al., 2016). Consistent with the immunostaining results using anti-CID, the CID-Dendra2 fusion displayed a 1.39-fold overall enrichment at sister chromatids segregating toward the future GSC side compared to the future GB side (Figures 1D and 1E, Movie S1). Additionally, CID-Dendra2 showed largely symmetric distribution patterns in SGs (Figures 1D and 1F, Movie S2). Finally, a CID-GFP genomic transgene used in previous studies (Henikoff et al., 2000) showed a 1.73-fold enrichment toward the future GSC side compared to the future GB side in asymmetrically dividing GSCs (Figures S1E and S1G, Movie S3). By contrast, a nearly equal distribution of CID-GFP was found in symmetrically dividing SGs (Figures S1F and S1G, Movie S4). In summary, both endogenous and transgenic CID showed enrichment toward the future stem cell side during GSC asymmetric division, using both immunostaining on fixed samples and live cell imaging.

We hypothesize that the asymmetries in CID segregation observed in anaphase and early telophase GSCs are due to the asymmetry in CID levels between individual pairs of sister centromeres. To test this hypothesis, we examined sister centromeres at prophase or prometaphase. In GSCs, sister centromeres that could be resolved in prophase and prometaphase already displayed quantitative differences: Sister centromeres had a 1.52-fold average asymmetry in CID levels (Figures 1G and 1I, Figures S1H and S1I). Conversely, the vast majority of sister centromeres that were resolved in prometaphase SGs showed no obvious level differences (Figures 1H and 1I). Even though a range of patterns could be detected in both GSCs and SGs (Figures S1H-J), the overall percentage of asymmetric sister centromeres in GSCs was significantly higher than that in SGs (Figure S1H). These results also suggest that the establishment of asymmetric CID at sister centromeres in GSCs likely occurred prior to prophase and prometaphase, when this asymmetry could be detected.

### The Temporal and Spatial Distribution of the CAL1 Chaperone in GSCs

In *Drosophila*, it has been demonstrated that Chromosome alignment defect 1 (CAL1) is both necessary (Erhardt et al., 2008; Goshima et al., 2007) and sufficient (Chen et al., 2014) to interact with CID and assemble CID-containing nucleosomes in the centromeric region. Using CRISPR/Cas9 technology, we generated a *Dendra2* knock-in line at the endogenous *cal1* gene. In non-replicative cells such as hub cells, CAL1-Dendra2 was undetectable (Figure S2A). In GSCs, centromeric localization of the CAL1-Dendra2 fusion protein was found prior to mitosis in G2 phase, as well as in early M phase (from prophase to prometaphase, Figure 2A). Even though CAL1 was detectable in mitotic GSCs, its level decreased as the cell progressed from G2 through early stages of mitosis (Figures 2A and 2B). However, at prophase or prometaphase when sister centromeres could be resolved in GSCs, asymmetric distribution of CAL1-Dendra2 could be detected, along with the asymmetric CID (Figures 2C and 2D, S2B and S2C). The percentages of asymmetric CAL1 (Figure S2B and S2C) is comparable with the percentages of asymmetric CID distribution at sister centromeres (Figures S1H and S1I).

**Figure 2:**
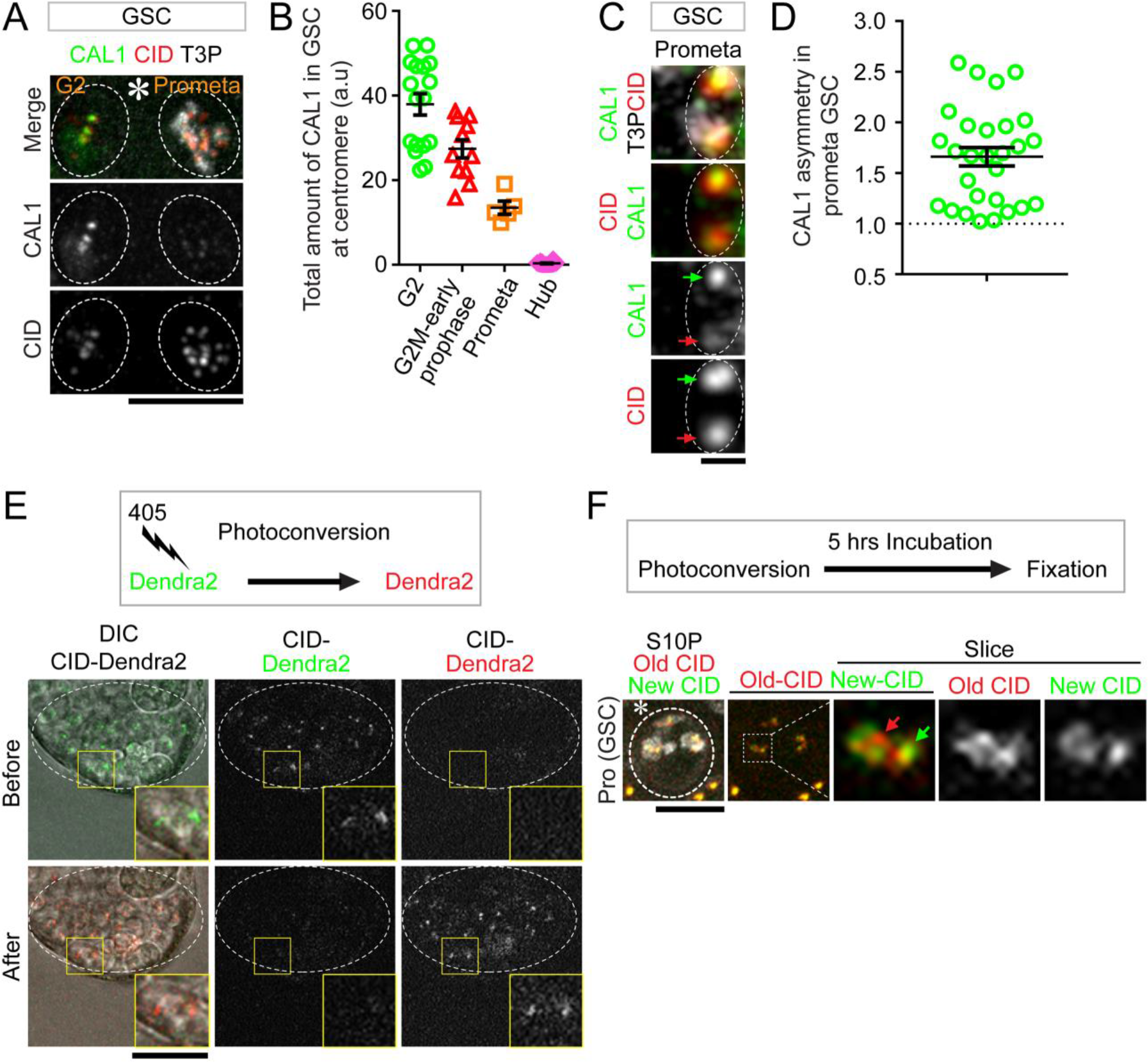
The temporal and spatial distribution of the CAL1 chaperone in male GSCs. (**A**) Two GSCs from one testis, one in G2 phase and the other in prometaphase, show higher CAL1 levels in G2 phase compared to prometaphase, visualized within the same testis using the same parameters. (**B**) Quantification of total amount of CAL1 in GSCs at different cell cycle stages: the highest CAL1 levels are detected in G2 phase, which gradually decreases in mitosis. The CAL1 signals in hub cells are used as a control. (**C**) In prometaphase GSCs, resolved individual sister centromeres show asymmetric distribution of CAL1 (green arrow-stronger centromere with more CID and CAL1, red arrow-weaker centromere with less CID and CAL1). (**D**) Quantification of individual pairs of resolved sister centromeres in prometaphase GSCs show asymmetric CAL1 distribution (1.66± 0.09-fold, *n* = 28), Table S5. (**E**) Photoconversion experiments at the apical tip of testis (circled area), which convert CID-Dendra2 protein from green fluorescent protein to red fluorescent protein. Insets show the very tip of the testis where GSCs are located. (**F**) After photoconversion, testes were cultured for 5hrs before fixation. New CID (green Dendra2) incorporation is detectable in early prophase GSCs. All ratios = Avg± SE, Asterisk: hub. Scale bars: 10μm (**A**), 0.5μm (**C**), 25μm (**E**), 5μm (**F**).

Unlike the canonical histones such as H3, deposition of the histone variant CID is not dependent on DNA replication, and the timing could be cell type-specific (Dunleavy et al., 2012; Garcia Del Arco et al., 2018; Mellone et al., 2011; Schuh et al., 2007). The asymmetry of CAL1-Dendra2 at sister centromeres indicates that it may deposit CID differentially onto the sister chromatids. To study the timing of new CID incorporation in *Drosophila* male GSCs, we utilized the knock-in *cid* gene that produces the CID-Dendra2 fusion protein. Dendra2 is a photoconvertable protein that can switch irreversibly from green fluorescence to red fluorescence. Following photoconversion any newly synthesized CID will be green fluorescent. Therefore, old CID is labeled in red while new CID labeled in green. We first checked that CID-Dendra2 had approximately 90% photoconversion efficiency in cells at the apical tip of testis (Figure 2E) (Experimental Procedures). To analyze the timing of new CID incorporation in GSCs, we photoconverted cells located at the testis tip and visualized mitotic GSCs five hours later (Figure 2F). Considering the elongated G2 phase in GSCs lasting for approximately 10-12 hours (Sheng and Matunis, 2011; Tran et al., 2012; Yadlapalli et al., 2011; Yadlapalli and Yamashita, 2013), the significantly increased green fluorescence detected in prophase GSCs indicate that new CID has been incorporated likely prior to mitosis in late G2 phase or in early mitosis (Figures 2F and S2D). As a control, the non-replicative hub cells showed no new CID incorporation during the same amount of time (Figure S2D), consistent with the lack of CAL1 expression in hub cells (Figure S2A).

In summary, we found that the CAL1 chaperone, which is responsible for deposition of the centromere-specific histone variant CID, has an asymmetric enrichment at sister centromeres in prophase and prometaphase GSCs when sister centromeres could be distinguished. This asymmetric distribution of CAL1 likely regulates new CID deposition during late G2 phase and possibly early M phase. We propose that the spatially- and temporally-specific CAL1 distribution and function are responsible for the asymmetric CID levels at sister centromeres.

### Asymmetric Kinetochore Correlates with the Asymmetry between Sister Centromeres in GSCs

The recognition of centromeres by microtubules is mediated by the kinetochore, a highly organized multiprotein structure that nucleates at the surface of centromere and coordinates the attachment of the mitotic spindle (Cheeseman, 2014; Cleveland et al., 2003). We next examined a key kinetochore protein, NDC80 (Schittenhelm et al., 2007), using a *Dendra2-Ndc80* knock-in fly strain. Consistent with the CID asymmetry, the kinetochore component NDC80 displayed a 1.49-fold overall enrichment at sister chromatids segregated toward the future GSC side in anaphase or early telophase GSCs (Figures 3A and 3C). In contrast, the symmetrically dividing SGs showed a nearly symmetric NDC80 segregation pattern (Figures 3B and 3C).

**Figure 3:**
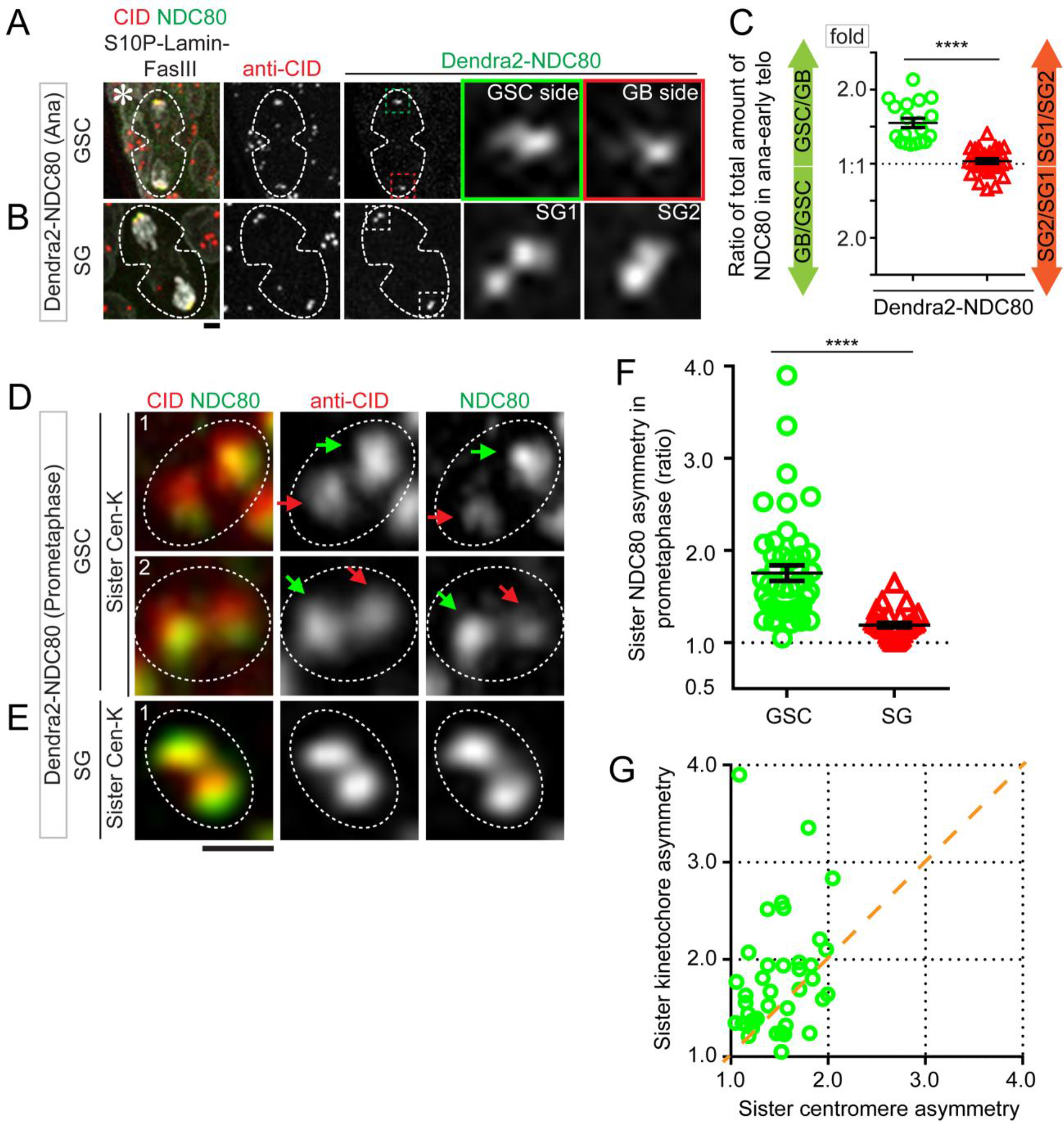
Asymmetric sister kinetochore correlates with the asymmetry of sister centromeres in GSCs. (**A**) Asymmetric kinetochore component NDC80 in asymmetrically dividing GSCs with more NDC80 toward the GSC side, using immunostaining of a Dendra2-NDC80 knock-in line. (**B**) In symmetric SG cell division, NDC80 is symmetrically distributed. (**C**) Quantification of both data sets: GSC/GB = 1.49± 0.07 (*n* = 19), SG1/SG2 = 1.03± 0.03 (*n* = 28), Table S6. (**D-E**) In prometaphase, resolved sister kinetochores show more asymmetry in GSCs (NDC80 in **D**, with the same polarity with CID enrichment), compared to resolved sister kinetochores in SGs (**E**). (**F**) Quantification of individual pairs of resolved sister kinetochores in prometaphase: 1.76± 0.08-fold in GSCs (*n* = 46); 1.19± 0.02-fold in SGs (*n* = 34), Table S7. (**G**) In prometaphase GSCs, asymmetric NDC80 is positively correlated with asymmetric CID. The degree of asymmetry for sister kinetochore is more than that for sister centromere [CID = 1.49-fold, NDC80 = 1.76-fold (*n* = 43), Table S8]. All ratios = Avg± SE. *P*-value: paired *t* test: ****: *P* < 10^−4^, Asterisk: hub. Scale bars: 2μm (**A-B**) and 0.5μm (**D-E**).

Furthermore, differences between individual pairs of sister kinetochores were already detectable in prometaphase GSCs (Figure 3D), but not in prometaphase SGs (Figure 3E). Individual pairs of sister kinetochores showed, on average, a 1.76-fold difference in prometaphase GSCs (Figure 3F, Figures S3A and S3B). Although a range of distinct patterns among sister kinetochores in prometaphase SGs could be detected, the asymmetry was considerably less compared to that in prometaphase GSCs (Figure 3F, Figures S3A and S3C).

Noticeably, sister kinetochore showed a similar asymmetric pattern compared to sister centromere (i.e. the side with more CID also had more NDC80, Figure 3D), but quantitatively sister kinetochores showed a greater degree of asymmetry than the sister centromeres (Figures 3D and 3G). Collectively, these results demonstrate that sister kinetochores are also quantitatively different in mitotic GSCs, differentially preparing sister chromatids for asymmetric attachment. Together, the asymmetric sister centromeres may serve as a *cis*-mechanism for differential recognition by sister kinetochores (Figure S3D).

### Temporally Asymmetric Microtubule Dynamics in GSCs

Since NDC80 is a key microtubule binding protein at the kinetochore (Cheeseman et al., 2006; DeLuca et al., 2006), we next sought to understand how asymmetry on sister kinetochores could be recognized by the mitotic spindle. We tracked microtubules using a GFP-tagged α-Tubulin under the control of an early-stage germline-specific *nanos-Gal4* driver (Van Doren et al., 1998), as used previously in this system (Yamashita et al., 2003). Using high temporal resolution movies, we tracked microtubules in real-time throughout the cell cycle of germ cells at distinct stages, including GSCs and SGs. Due to the unique morphology of the mitotic spindle at metaphase, we used metaphase as a landmark to define time point zero; other cell cycle phases before metaphase were labeled accordingly. In male GSCs, the mother centrosome is retained in proximal to the GSC-niche interface while the daughter centrosome migrates to the distal side after duplication in early G2 phase (Yamashita et al., 2003; Yamashita et al., 2007). From −250 minutes (−250’) at approximately mid-G2 phase to −150’ at late-G2 phase, more microtubules emanated from the mother centrosome than microtubules from the daughter centrosome (Figure 4A, Movie S5). Subsequently, the dominance of mother centrosome-emanating microtubules declined at the G2-to-M phase transition from −50’ to −35’, as the daughter centrosome-emanating microtubules increased (Figures 4A and 4C, Movie S5). This decrease of microtubules from the mother centrosome and increase of microtubules from the daughter centrosome persisted from −30’ to −10’ in prophase GSCs, as well as at −5’ in prometaphase GSCs (Figures 4A and 4C, Movie S5). At metaphase, microtubules from both centrosomes were almost equal, forming the mitotic spindle perpendicular to the GSC-niche interface (Figures 4A and 4C, Movie S5), as reported previously (Yamashita et al., 2003).

**Figure 4:**
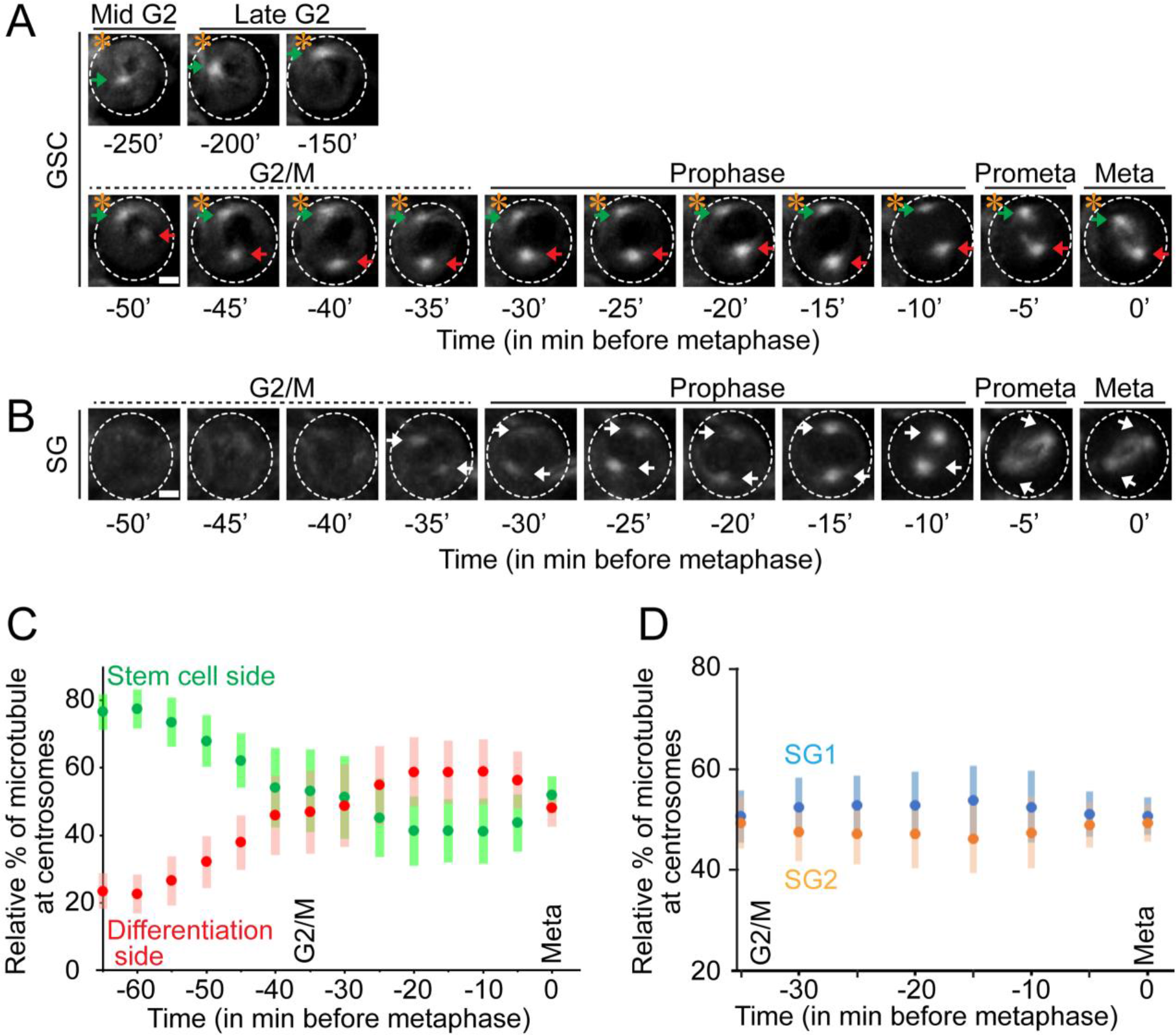
Temporally asymmetric microtubules in GSCs. 3D reconstructed montage made from live cell imaging using a *nanos-Gal4*; *UAS-α-tubulin-GFP* line in GSCs (**A**) and quantified in (**C**, *n* = 22, Table S8), as well as in SGs (**B**) and quantified in (**D**, *n* = 24, Table S9). Metaphase was set as time 0, and other time points prior to metaphase were labeled as minus minutes accordingly. (**A**) In GSCs the mother centrosome is activated in mid G2 (−250 min) prior to the daughter centrosome at G2/M transition (−50 min). (**B**) In SGs, both centrosomes are active at the G2/M transition (−35 min) simultaneously. (**C-D**) All ratios = Avg± SE, see Table S8. Asterisk: hub. Scale bars: 2μm.

Conversely, in SGs, microtubules from both centrosomes became simultaneously detectable at −35’ at the G2-to-M phase transition. After that, no obvious difference in microtubules could be detected between the two centrosomes throughout mitosis (Figures 4B and 4D, Movie S6). In summary, these results demonstrate that microtubules, the key component of the mitotic machinery, display a temporal asymmetry in male GSCs compared to SGs.

### Polarized Nuclear Envelope Breakdown Induced by Temporally Asymmetric Microtubules

Next, using high spatial resolution movies, we detected that microtubules emanating from the mother centrosome were able to physically dissemble and penetrate the nuclear envelope from late G2 phase to early prophase in GSCs (Figure S4A, Movie S7). Using antibodies against *Drosophila* Lamin B (Chen et al., 2013), we found that the dynamic activity of microtubules derived from the mother centrosome resulted in the invagination of the nuclear lamina exclusively at future stem cell side at early prophase (Figure 5A, Movie S8A-C). Local disassembly of nuclear lamina was first detected at the future stem cell side (green arrows in Figure 5A and Movie S8A-C), after which increased microtubule dynamics from the daughter centrosome led to invagination of nuclear lamina from the GB side, leading to nuclear envelope breakdown (NEBD, red arrowheads in Figure 5B and Movie S9A-B). By contrast, such a temporal polarity in NEBD was not observed in symmetrically dividing SGs (yellow arrowheads in Figure 5C and yellow arrows in Figure 5D).

**Figure 5:**
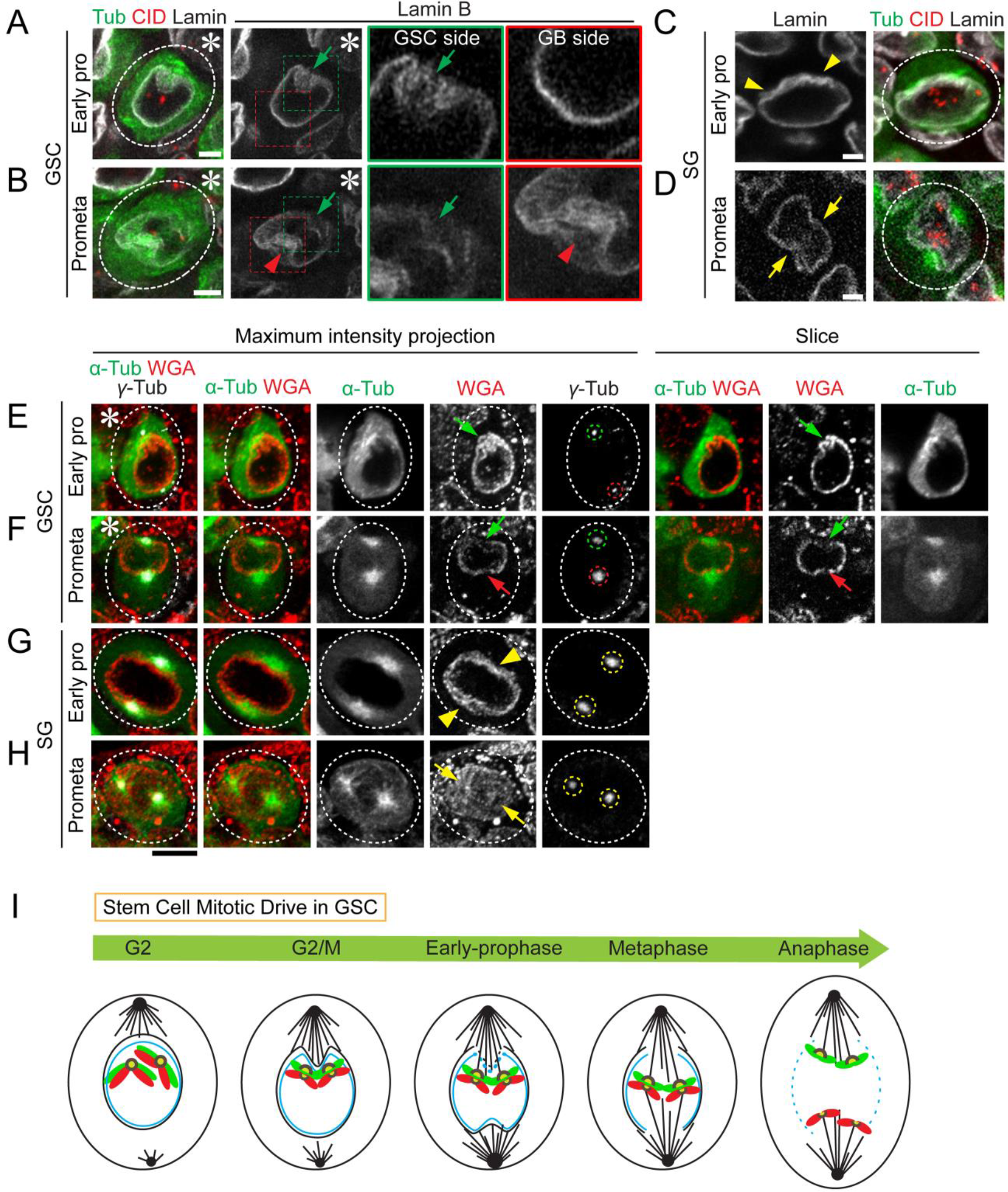
Polarized NEBD in GSCs. (**A-B**) Morphology of nuclear envelope in GSCs at early prophase (**A**) and at prometaphase (**B**), as well as in SGs at early prophase (**C**) and at prometaphase (**D**), visualized by immunostaining using anti-Lamin B (white), co-stained with anti-CID (red) using *nanos-Gal4*; *UAS-α-tubulin-GFP* line (α-Tub in green). Early active mother centrosome-nucleating microtubules correspond with the initial NEBD toward the hub (green arrows in **A** and **B**), whereas later daughter centrosome-nucleating microtubules lead to NEBD toward the other side (red arrowhead in **B**). In comparison, SGs show no polarized NEBD (yellow arrowheads label invagination sites in **C**, while yellow arrows point to symmetric NEBD sites in **D**). (**E-H**) Morphology of nuclear envelope in GSCs (**E-F**) and SGs (**G-H**) at different cell cycle phases, visualized by wheat germ agglutinin (WGA, red) which binds to the cytoplasmic part of each nuclear pore, co-stained with anti-γ-Tubulin (white) in *nanos-Gal4*; *UAS-α-tubulin-GFP* line (α-Tub in green). The ‘poking in’ activities of microtubules are labeled by green arrows (from GSC side) and red arrows (from GB side) in (**E-F**). The interactions between microtubules and nuclear envelope in SGs are shown by yellow arrowheads at ‘poking in’ sites in (**G**) and yellow arrows at NEBD sites in (**H**). **(I)** A cartoon depicting the sequential events in mitotic drive in *Drosophila* male GSC. Asterisk: hub. Scale bars: 2 μm.

Similar results were obtained using wheat germ agglutinin (WGA), which binds to the cytoplasmic part of each nuclear pore, allowing for visualization of nuclear membrane morphology in GSCs (Figures 5E and 5F) and SGs (Figures 5G and 5H). Therefore, in GSCs the temporally asymmetric microtubule dynamics likely resulted in a polarized NEBD, first at the GSC side and then at the GB side.

We next asked whether the asymmetric activity of microtubules and polarized NEBD coordinates with chromosomes at the centromeric region, visualized by CID signals. Noticeably, in late G2 phase GSCs, close proximity between chromosomes and microtubules could be detected: When the mother centrosome-emanating microtubules dynamically interacted with the nuclear envelope (α-Tubulin in Figures S4B and S5A), centromeres from all chromosomes clustered preferentially toward the future GSC side (CID in Figures S4B and S5A). Given the higher microtubule dynamics from the mother centrosome, this close proximity of all centromeres toward the future GSC side might initiate crosstalk between centromeres and mother centrosome-emanating microtubules. A series of immunostaining images that showed microtubules (α-Tubulin), nuclear lamina (Lamin), the mitotic chromosome marker H3S10P (S10P), and centromeres (CID) illuminated coordinated microtubule activity and NEBD with chromosome and centromere movement in GSCs at different cell cycle stages (Figures S5A-F). As a comparison, we also examined these events in symmetrically dividing SGs: Even though microtubules emanating from both centrosomes still coincided with chromosomal condensation and centromere movement, no obvious sequential order was observed (Figures S5G-K), and no detectable asymmetry was found between resolved individual pairs of sister centromeres (CID in Figure S5H) or at anaphase between the segregated two sets of sister chromatids (CID in Figure S5K).

Together, these immunostaining images with markers for both the *trans*-factors (e.g. microtubule and nuclear lamina) and the *cis*-factors (e.g. CID) delineate a series of events at sister centromeres, allowing for differential recognition and segregation of sister chromatids (Figure S5L). This series of interactions has cellular specificity to asymmetrically dividing GSCs but not to symmetrically dividing SGs.

### Compromising the CAL1 Chaperone Abolishes the *Cis*-asymmetry of CID and Leads to Stem Cell Defects

Based on these results, we propose a two-step model for the establishment and recognition of CID asymmetry in GSCs (Figure 6A): Step 1 involves the establishment of asymmetric CID at individual pairs of sister centromeres from late G2 phase to early M phase; Step 2 uses asymmetric mitotic spindle activity to recognize asymmetric sister centromeres for differential attachment and segregation during ACD.

**Figure 6:**
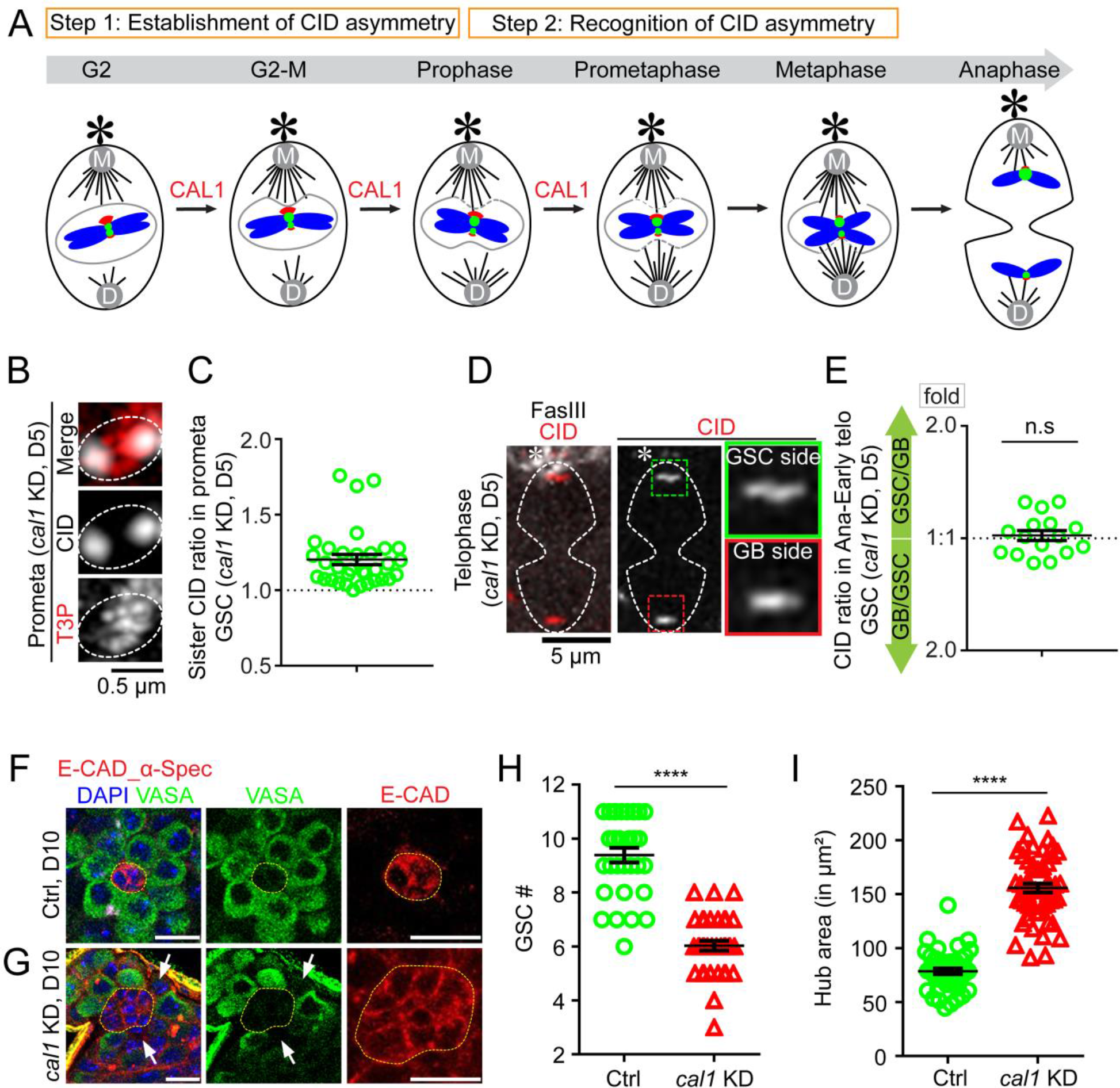
Compromising the CAL1 chaperone disrupts the CID asymmetry and results in GSC defects. (**A**) A cartoon depicting a two-step model for asymmetric CID inheritance: Step 1 involves establishment of CID asymmetry at individual sister centromeres; Step 2 requires recognition of asymmetric CID by the mitotic machinery. CAL1 is likely contributing to Step 1. (**B**) In prometaphase GSCs after knocking down *cal1* for five days (D5), resolved individual sister centromeres show symmetry, quantified in (**C**): 1.20± 0.03-fold, *n* = 33, Table S10. (**D**) Symmetric CID segregation in anaphase or early telophase GSCs after knocking down *cal1* for five days (D5), quantified in (**E**): 1.03± 0.03-fold, *n* = 16, Table S11. (**F-G**) Apical tip of testes from *tub-Gal80^ts^, nos-Gal4* (*Ctrl*, **F**) and *tub-Gal80^ts^, nos-Gal4 > cal1 shRNA* (*cal1* KD, **G**) males after knocking down *cal1* for ten days (D10), immunostained with a germ cell marker VASA (green), a hub marker *Drosophila* E-Cadherin (E-CAD, red), and a spectrosome/fusome marker α-Spectrin (α-Spec, red). The empty niche place without GSCs is labeled by white arrows in (**G**), the number of GSCs was quantified in (**H**): *Ctrl* (9.44± 0.25, *n* = 34), *cal1* KD (6.03± 0.19, *n* = 35), Table S12; hub area is indicated by yellow dotted outline in (**F**-**G**) and quantified in (**I**): *Ctrl* (80.44± 2.65μm^2^, *n* = 45), *cal1* KD (155.88± 4.05μm^2^, *n* = 55), Table S15. All ratios = Avg± SE. *P*-value: paired *t* test. ****: *P* < 10^−4^; n.s: not significant. Asterisk: hub. Scale bars: 0.5μm (**B**), 5μm (**D**), 10μm (**F**-**G**).

To investigate whether these two steps are important for GSC identity and activity, we next designed experiments to disrupt either Step 1 or Step 2. Because the CAL1 chaperone is ubiquitously required for CID incorporation, we compromised CAL1 in a spatiotemporally controlled manner to avoid any potential secondary effects. The temperature-sensitive Gal80 controlled by the *tubulin* promoter (*tub-Gal80^ts^*) was used in combination with the early germline-specific *nos-Gal4* driver, in order to temporally control the expression of *cal1 shRNA* (short hairpin RNA) in adult testes [*tub-Gal80^ts^, nos-Gal4 > cal1 shRNA* (*cal1* KD)]. We found that the CID asymmetry in anaphase or early telophase GSCs was still detectable two days after shifting from the permissive to the restrictive temperature as adult flies (D2). By contrast, at five days (D5) this asymmetry became almost undetectable (Figure S6A). At D5, the CAL1 protein was almost undetectable using anti-CAL1 for immunostaining (Figure S6B), suggesting an efficient knocking down.

We then examined whether *cal1* KD disrupts the establishment of CID asymmetry at individual sister centromeres in Step 1. Indeed, in D5 prometaphase GSCs, approximately 63.6% of sister centromeres displayed symmetric CID distribution compared to 15.4% in WT GSCs (Figures 6B and 6C, S6C-D *versus* S1H-I), suggesting that the establishment of CID asymmetry was disrupted in *cal1* KD GSCs. Not surprisingly, the asymmetric inheritance pattern of CID was lost in anaphase or early telophase GSCs from D5 males (Figures 6D and 6E), consistent with the prediction that the disruption of Step 1 would lead to symmetric inheritance patterns (Figure 6A).

Next, we asked if the loss of CID asymmetry could lead to any germ cell defects. We found that in *cal1* KD testes at D10 when the asymmetric CID segregation pattern was abolished for several days (Figure S6A), there was a significant loss of GSCs compared to the *tub-Gal80^ts^, nos-Gal4* (*Ctrl*) testes under the same condition (Figures 6F *versus* 6G, and 6H). However, the mitotic index was comparable between *cal1* KD GSCs and the *Ctrl* GSCs [*cal1* KD: 10.1% (48 H3S10P-positive GSCs/476 total GSCs) *versus Ctrl*: 13.1% (42 H3S10P-positive GSCs/321 total GSCs)], suggesting that the GSC loss phenotype upon knocking down CAL1 is not a result of compromised GSC proliferation but likely due to GSC maintenance defects. Finally, a dramatic increase of the hub area was detected in *cal1* KD testes compared to the *Ctrl* testes (hub region in Figure 6F *versus* 6G, and 6I), which should be a secondary effect due to GSC loss, as reported previously (Dinardo et al., 2011; Gonczy and DiNardo, 1996; Monk et al., 2010; Tarayrah et al., 2015; Tazuke et al., 2002; Xie et al., 2015). In summary, these results demonstrate that knockdown of the CAL1 chaperone in adult testes leads to symmetric CID distribution at individual sister centromeres followed by an overall symmetric segregation of CID in GSCs, as well as significant GSC loss and hub enlargement phenotypes (Figure S6E).

### Depolymerizing Microtubules Disrupts the *Trans*-asymmetry of the Mitotic Machinery and Results in Randomized Sister Chromatid Segregation

In order to investigate how temporal asymmetries in microtubules may regulate asymmetric sister chromatid attachment and segregation in the mitotic GSCs, we utilized the microtubule depolymerizing drug Nocodazole (NZ) to acutely destabilize microtubules in a temporally controlled and reversible manner. We applied NZ on testes *ex vivo* followed by washing out and immunostaining (Figure 7A) (Experimental Procedures). Here, NZ treated GSCs were arrested at the G2/M transition, but following washing out, they were released and progressed through mitosis in a time-dependent manner (Figures 7A and S7A).

**Figure 7:**
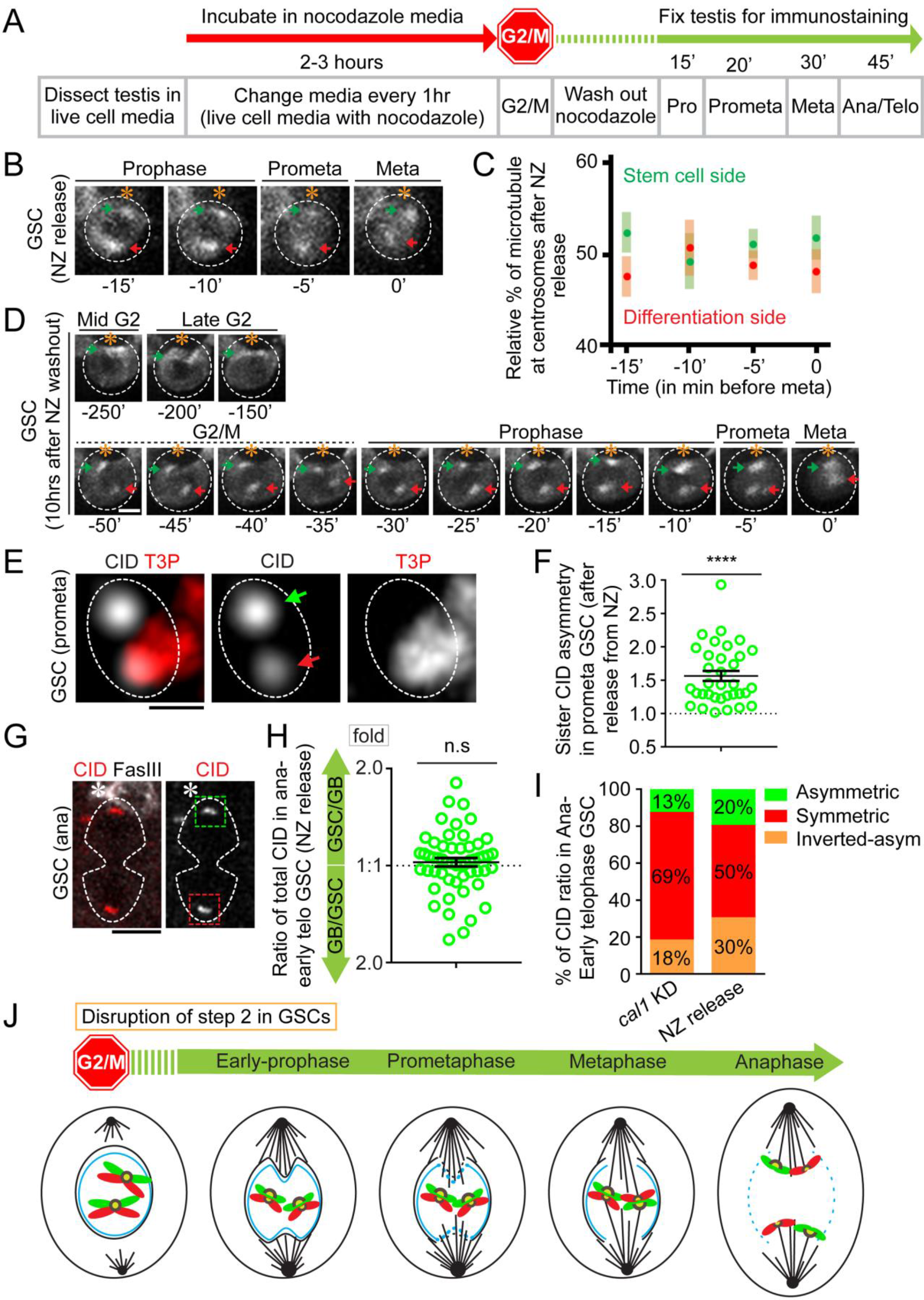
Depolymerizing microtubules disrupts the temporal microtubule asymmetry and results in randomized sister chromatid segregation. (**A**) A regime of Nocodazole (NZ) treatment experiment whereby testes were treated with NZ for two to three hours to arrest GSCs primarily at the G2/M transition. Immediately after washing out NZ, GSCs progress into different stages of mitosis in a time-dependent manner. (**B**) Immediately after releasing from NZ treatment, microtubules nucleate from both mother centrosome and daughter centrosome almost simultaneously, visualized by α-Tubulin-GFP (green) with 3D reconstructed montage made from live cell imaging. (**C**) Quantification of percentage of microtubules emanated from mother centrosome *versus* microtubules emanated from daughter centrosome in GSCs (*n* = 10, Table S14). (**D**) Recovery of asymmetric microtubules emanated from mother centrosome (green arrow) *versus* daughter centrosome (red arrow) in GSCs approximately 10hrs after releasing from NZ treatment. (**E**) In prometaphase GSCs immediately after releasing from NZ treatment, resolved sister centromeres display asymmetric distribution (green arrow: stronger centromere, red arrow: weaker centromere). (**F**) Quantification of individual pairs of resolved sister centromeres in prometaphase GSCs immediately after releasing from NZ treatment: 1.56± 0.08-fold (*n* = 34), Table S15. **(G)** Randomized CID segregation pattern in anaphase or early telophase GSCs immediately after releasing from NZ treatment, quantified in (**H**): 1.05± 0.03-fold, *n* = 54, Table S16. (**I**) Percentage of GSCs showing conventional asymmetric (see Supplemental Information for classification, GSC side/GB side ≥ 1.2), symmetric (0.9 < GSC side/GB side < 1.2), and inverted asymmetric (GSC side/GB side ≤ 0.9, i.e. more toward GB side than to the GSC side) patterns of CID segregation at anaphase or early telophase, in *cal1* KD (disrupts Step 1) and immediate release from NZ treatment (NZ release, disrupts Step 2). More symmetric segregation patterns are seen in *cal1* KD compared to NZ release; both conventional asymmetric and inverted asymmetric patterns are detected with NZ release, suggesting the segregation pattern is randomized. (**J**) A cartoon depicting randomized sister chromatids segregation upon disrupting Step 2 without affecting Step 1. Asterisk: hub. All ratios = Avg± SE. *P*-value: paired *t* test. ****: *P* < 10^−4^; n.s: not significant. Scale bars: 5μm (**B, D, G**), 0.5μm (**E**).

Immediately after releasing from NZ-induced arrest, the temporal asymmetry of microtubules was compromised, as both the mother centrosome and the daughter centrosome showed comparable microtubule-emanating activity (Figures 7B and 7C, Movie S10). Furthermore, the polarity in NEBD detected in untreated GSCs (Figures 5A to 5B and S5B to S5E) was disrupted soon after releasing from NZ treatment (α-Tubulin and Lamin in Figures S7B-C). These results demonstrate that NZ arrest and release disrupts temporal asymmetries in microtubule activity as well as the downstream effects on nuclear envelope disassembly. We found that these effects were acute, as GSCs were able to recover asymmetric microtubule activity following a 10-hour recovery period after washing out NZ (Figure 7D, Movie S11). Noticeably, in GSCs, the centrosomes were already duplicated and well oriented perpendicular to the GSC-niche interface at early prophase (α-Tubulin in Figure S7B), whereas in SGs duplicated centrosomes were in proximity with each other at early prophase (α-Tubulin in Figure S7D) and were at the opposite sides only later during mitosis (α-Tubulin signals in Figure S7E). This observation is consistent with a previous report that revealed centrosomes in GSCs duplicate and migrate during the G2 phase and prior to mitosis (Yamashita et al., 2003). This observation also confirmed that the NZ treatment followed by release changed microtubule emanation in an acute manner, since the centrosome duplication and migration were not disrupted. Taken together, these results suggest that by arresting and releasing GSC with the microtubule depolymerizing drug NZ, we are able to selectively disrupt the temporal asymmetry of microtubule activity in GSCs, which allow us to assess the impacts of loss of microtubule asymmetry in GSCs.

We next examined CID distribution at individual sister centromeres in prometaphase GSCs and found that this asymmetry was largely maintained (Figures 7E and 7F, S7F and S7G). These results demonstrate that acute disruption of microtubule activity is not required for establishment of asymmetric sister centromeres. However, asymmetry between individual pairs of sister kinetochores was greatly reduced (Figure S7H), suggesting that asymmetries in microtubule activity may help establish sister kinetochore asymmetries. Additionally, under this condition, asymmetric CID segregation in anaphase and early telophase GSCs was lost (Figures 7G and 7H). Overall, GSCs after NZ release showed a significantly higher incidence of symmetric CID segregation pattern, as well as both asymmetric (more CID toward GSC side than to GB side) and inverted asymmetric (more CID toward GB side than to GSC side) patterns (Figures 7H and 7I). Both asymmetric ratios were greater than those in *cal1* KD GSCs (Figure 7I). These results further confirmed that Step 2 was disrupted without affecting Step 1 using this regime (Figure 6A): When Step 1 was disrupted in *cal1* KD GSCs, most GSCs displayed a symmetric CID segregation patterns. However, when Step 2 was disrupted without affecting Step 1, randomized CID segregation patterns were detectable including both asymmetric and inverted asymmetric patterns (Figure 7I). Moreover, asymmetric sister centromeres with a randomized orientation could be observed in metaphase GSCs soon after NZ release (Figures S7I and S7J). Together, these results demonstrate that while temporal asymmetries in microtubule activity are dispensable for establishment of asymmetric sister centromeres, these asymmetries in microtubule activity are essential for the recognition and coordinated segregation of asymmetric sister chromatids (Figure 7J).

Consistent with our hypothesis that NZ treatment randomizes sister chromatid inheritance, both old and new histone H3 displayed randomized segregation patterns at anaphase or telophase GSCs immediately following NZ release (Figures S7K and S7L). Furthermore, upon recovery after releasing from NZ treatment, some GSCs could restore the old H3 asymmetry (Figure S7M), consistent with the acuteness of this treatment. Taken together, randomized CID and H3 inheritance patterns were detected as a result of immediate release from NZ-induced arrest. These results suggest that the temporal asymmetries from the mitotic machinery are required for recognizing and segregating the asymmetric sister centromeres during the process of asymmetric histone inheritance (Figure S7N).

## Discussion

Here, we have shown that asymmetrically dividing GSCs utilize a cohort of *trans-* and *cis-* factors to recognize and segregate epigenetically distinct sister chromatids. The previously identified asymmetrically inherited centrosomes (Yamashita et al., 2007), the herein reported temporally asymmetric microtubules, the polarized NEBD, and the asymmetric sister kinetochores serve as an axis of asymmetric *trans-*factors within the mitotic machinery. Meanwhile, the previously identified asymmetric H3 (Tran et al., 2012), the differential phosphorylation at Thr3 of H3 (Xie et al., 2015), and the herein reported asymmetric sister centromeres act as a series of *cis-*factors. Together, temporally asymmetric microtubules coordinate with asymmetric sister centromeres to ensure differential sister chromatid attachment followed by segregation to generate two epigenetically distinct daughter cells, a phenomenon that is consistent with what has been previously hypothesized (Kahney et al., 2017; Malik, 2009; Yamashita, 2013). We propose that this series of interactions represent a ‘mitotic drive’ phenomenon during the asymmetric division of *Drosophila* male GSCs (Figure 5I).

Mitotic drive requires that both *cis*- and *trans*-factors work together to ensure asymmetric sister chromatid segregation (two steps in Figure 6A). In order to understand how key aspects of this process are regulated, we designed several experiments to investigate both the establishment of the *cis*-asymmetry (Step 1) and the recognition by the *trans*-factors (Step 2). First, using knockdown experiments we demonstrated that establishment of the CID asymmetry requires the activity of CAL1 chaperone. When Step 1 was disrupted using the *cal1* knockdown condition, individual pairs of sister centromeres show symmetry and the symmetric segregation pattern is prevalent in anaphase or telophase GSCs (Figures 6B-E and 7I). We then developed a drug arrest and release protocol to test whether recognition of the CID asymmetry depends on the mitotic machinery. Our results demonstrate that in the absence of temporally asymmetric microtubule activity, asymmetries between individual pairs of sister centromeres are still preserved through the actions of the CAL1 chaperone (Figures 7E-F and S7F-G). However, loss of temporal asymmetry in mitotic machinery severely compromised the proper recognition and segregation of asymmetric sister centromeres, leading to a range of CID segregation patterns from asymmetric, inverted asymmetric to symmetric patterns (Figure 7H-I). Because male flies have two sex chromosomes (X and Y chromosomes) and two major autosomes (2^nd^ and 3^rd^ chromosomes), even randomized segregation of sister chromatids could lead to an asymmetric pattern, albeit at a low percentage as shown previously (Xie et al., 2015). In theory, the low occurrence of the asymmetric patterns could have the opposite polarities with comparable ratios, which was what we observed in anaphase or telophase GSCs immediately after releasing from NZ treatment (Figure 7I). These results demonstrate that these two steps could be uncoupled and disruption of either step could lead to largely symmetric segregation pattern of sister chromatids at anaphase or telophase, but distinct patterns at individual pairs of sister centromeres at prophase or prometaphase.

In general, ACD gives rise to two daughter cells with distinct fates, which could occur during development, as well as tissue homeostasis and regeneration (Clevers, 2005; Kahney et al., 2017; Knoblich, 2010; Morrison and Kimble, 2006; Venkei and Yamashita, 2018). Many studies have helped our understanding how this difference in cell fate could be achieved in stem cells. While extrinsic cues contribute to ACD, asymmetrically segregated intrinsic factors could also be critical for cell fate decisions. More than four decades ago, an “immortal strand” hypothesis proposed that stem cells retain “immortal” DNA strands (Cairns, 1975) to minimize the incidence of mutations, which may arise during pathological progression or aging. However, later *in vivo* studies using more rigorous methods and precise labeling have challenged this hypothesis (Klar and Bonaduce, 2013; Sauer et al., 2013; Schepers et al., 2011; Yadlapalli and Yamashita, 2013). On the other hand, examples of biased chromosome segregation have been shown in multiple systems, including both invertebrates and vertebrates (Rocheteau et al., 2012; Yadlapalli and Yamashita, 2013). It has been proposed that epigenetic differences between sister chromatids, especially at the centromeric region, are required to direct the asymmetric outcomes during ACD (Klar, 1994, 2007; Lansdorp, 2007). However, no *in vivo* evidence in any multicellular organism has been shown to support these hypotheses. It is conceivable that the centromere and kinetochore asymmetry may provide cellular mechanisms for selective sister chromatid segregation during ACD (Yadlapalli and Yamashita, 2013). Interestingly, asymmetric kinetochores were reported previously as a mechanism to define a specific lineage in yeast *Saccharomyces cerevisiae* (Thorpe et al., 2009). Yet, it remains unclear whether a similar phenomenon could occur in multicellular organisms. Here, our findings not only provide the first direct *in vivo* evidence in support of these hypotheses, but also reveal the cellular and molecular specificities of such selective chromatid segregation in a well characterized adult stem cell model.

The “meiotic drive” hypothesis has proposed that the allele with a stronger kinetochore is retained in the oocyte while the allele with a weaker kinetochore is segregated to polar bodies for degeneration during female meiosis (Henikoff et al., 2001; Malik and Henikoff, 2009; Pardo-Manuel de Villena and Sapienza, 2001a, b). It has been shown that in female mice, the stronger kinetochore often associates with the longer, “selfish” centromere and has more affinity to the meiotic spindle. Moreover, the meiotic spindle itself has been asymmetrically modified due to polarized signaling (Akera et al., 2017; Chmatal et al., 2014; Iwata-Otsubo et al., 2017; Kursel and Malik, 2018). Recently, it has also been shown that a microtubule motor protein regulates the meiotic drive in maize, indicating a role for microtubules in selective attachment to centromeres in this process (Dawe et al., 2018; Schroeder and Malik, 2018).

Despite these exciting new discoveries in the field of meiosis, it is unclear whether these asymmetric functions of the kinetochore and centromere could act in mitosis to regulate sister chromatid segregation. Here, our results reveal cellular and molecular mechanisms in mitosis that have both commonalities and distinctions compared to meiotic drive. First, in meiosis, the centromere difference occurs between specific homologous chromosomes, whereas in mitosis, it occurs between genetically identical sister chromatids. Second, in meiosis, the microtubule itself does not have temporally asymmetric activity, but has different modifications instead. In contrast, in mitosis temporal differences in microtubule nucleation activity from both centrosomes could be detected, which might bias attachment of microtubules to epigenetically distinct centromeres. Third, the biological outcome is different: The meiotic drive leads to the retention of a particular allele with a stronger kinetochore and a “selfish” centromere to the haploid oocyte, whereas the mitotic drive leads to two diploid cells, each with distinct epigenetic information for distinct cell fates. Although the meiotic drive and the stem cell mitotic drive have distinct features, they also share certain commonalities at the cellular level. For example, in both processes, the centromere difference serves as a key *cis*-acting mechanism while microtubules act as the crucial *trans*-factor to recognize the centromeric difference, which leads to differential inheritance of chromosomes.

In summary, our results demonstrate that the *cis*-asymmetry in sister chromatids tightly coincides and coordinates with the *trans*-asymmetry from the mitotic machinery. Together, this spatiotemporally-regulated axis of asymmetry, i.e. sister centromeres → sister kinetochores → nuclear membrane→ microtubules, allows for differential attachment and segregation of epigenetically distinct sister chromatids. Together, our discovery helps further understanding of a fundamental biological question regarding how the mitotic machinery could distinguish and ensure asymmetric partitioning of epigenetic information. This work also opens a new avenue to study whether and how similar mechanisms may be used in other contexts of multicellular organisms to generate cells with distinct fates.

## Acknowledgements

We thank Y. Yamashita, S. Erhardt, M. Van Doren, R. Johnston and X.C. lab members for insightful suggestions. We thank J. Gall for WGA, S. Erhardt for anti-CID, R. Quinn for technical assistance, Johns Hopkins Integrated Imaging Center for confocal imaging. Supported by NIGMS/NIH T32GM007231 (J.S.), NIGMS/NIH R01GM112008 and R35GM127075, the Howard Hughes Medical Institute, the David and Lucile Packard Foundation, and Johns Hopkins University startup funds (X.C.).

## Author contributions

R.R., J.S. and X.C. conceptualized the study. R.R. and J.S. performed all the experiments and data analysis. R.R. and X.C. wrote the manuscript.

## Competing interests

The authors declare no competing interests.

## Materials & Correspondence

Correspondence and material requests should be addressed to X.C.

