## Supplemental Information for "Stem cell mitotic drive ensures asymmetric epigenetic inheritance"

### Supplemental Experimental Procedures

#### Fly strains and husbandry

Fly stocks were raised using standard Bloomington medium at 18°C, 25°C, or 29°C as noted. The following fly stocks were used: *hs-flp* on the X chromosome (Bloomington Stock Center BL-26902), *nos-Gal4* on the 2nd chromosome (Van Doren et al., 1998), *UASp-FRT-H3-GFP-PolyA-FRT-H3-mKO* on the 3rd chromosome as reported previously (Tran et al., 2012), *UASp-FRT-H3-mCherry -PolyA-FRT-H3-eGFP* on the 3rd chromosome, *UAS-α-Tubulin-GFP (*Bloomington Stock Center BDSC #7373*)* on the 3rd chromosome, *GFP-cid (*Bloomington Stock Center BDSC #25047*)* on the 2nd chromosome, *nos-Gal4 (without VP16)/Cyo; tub-Gal80$^{ts}$/TM6B* (from Yukiko Yamashita, University of Michigan, Ann Arbor, Michigan, USA), and *UAS-cal1-shRNA* (BDSC # 41716) on the 3rd chromosome.

#### Generating knock-in fly strains

In collaboration with Fungene Inc. (Beijing, China), the following fly lines were generated using the CRISPR-Cas9 technology: CG613329 (*cid*) with Dendra2 tag at the internal site (between 118th - 119th codon), in order to generate the following fusion protein: CID N term-Dendra2-CID C term; CG5148 (*cal1*) with Dendra2 tag at the 5' immediately downstream of the START codon, in order to generate the following fusion protein: Dendra2-CAL1; CG9938 (*Ndc80*) with Dendra2 tag at the 5' immediately downstream of the START codon, in order to generate the following fusion protein: Dendra2-NDC80.

#### Photoconversion of Dendra2 protein

*Drosophila* testes from CID-Dendra2 knock-in strain were dissected and mounted in the live cell medium in a fluorodish, and put under the LSM 780 microscope. The 40X objective was used, the region of interest at the apical tip of the testis was selected; the 405 laser at 10-15% laser power for 30sec/pulse and ~100 iterations were used to photoconvert Dendra2 protein from green fluorescent protein to red fluorescent protein. The photoconversion efficiency was measured by quantifying amount of green Dendra2 before photoconversion (Db) and the amount of green Dendra2 immediately after photoconversion (Da) using the following formula: Photoconversion efficiency= (Db-Da)/Db X 100 %.

**Immunostaining**

Immunofluorescence staining was performed using standard procedures (Hime et al., 1996; Tran et al., 2012). The primary antibodies used were mouse anti-Fas III (1:100, DSHB, 7G10), rabbit anti-Vasa (1:200, Santa Cruz, Cat# sc-30,210), mouse anti α-Spectrin (1:50, DSHB, Cat# 3A9), mouse anti-Armadillo (1;200, DSHB, Cat# N27A1), rat anti-DE-CAD (1;50, DSHB, Cat# DCAD2), rat anti-CID (1:200, Active motif, Cat# 61735), rabbit anti-CID (1:200, Active motif, Cat# 69719), mouse anti-Lamin B (1:200, DSHB, Cat# ADL67.10), mouse anti-H3S10P (1:2,000, Abcam, Cat# ab14955), and rabbit anti-H3T3P (1:200, Millipore, Cat# 05-746R). Secondary antibodies were the Alexa Fluor-conjugated series (1:1,000; Molecular Probes). Images were taken using Zeiss LSM 700 confocal microscope, Zeiss LSM 780 confocal microscope, or Zeiss LSM 800 confocal microscope with Airyscan with 63x oil immersion objectives. Airyscan processing was performed to resolve sister centromeres (Figures 1G-I, 2C-D, 3D-G, 6B-C, 7E-F, S1I-J, S2B-C, S3B-C, S5C, S5H, S6C-D, and S7F-J), to detect microtubules (Figures S4A and S7I), and polarized NEBD (Figures S4A, 5A-D, and S5A-K).

Images were processed using Imaris software (3D image reconstruction) and Fiji software (to quantify the total amount of protein "RawIntDen") or to generate maximum intensity projection.

**Chromosome spreading**

Chromosome spreading was performed to better visualize sister centromeres and sister kinetochores at prophase and prometaphase. Adult *Drosophila* testes were dissected in Schneider's medium and incubated in a hypotonic solution (0.5% sodium citrate) for 5 minutes (min) at room temperature (RT). Testes were then fixed in freshly prepared 4 % paraformaldehyde in PBS for 5-7 min at RT on a Superfrost plus slide, a cover slip was placed on top of fixed tissue, and squashed as hard as possible with a thumb. Testes were then frozen by immersing them in liquid nitrogen, the cover slip was popped off using a razor blade, followed by immediate incubation in chilled 95% ethanol (-20°C) for 10 min. The immobilized testes were washed three times with 1x PBST, 10 min each time, followed by overnight incubation with primary antibodies in blocking solution (1x PBST with 3% BSA) at 4°C. The testes were then washed three times with 1x PBST, 15 min each time, followed by incubation with secondary antibodies for two hours at RT, washed three times with 1x PBST, 15 min each time, and mounted in Vectashield without DAPI (Vector, Cat# H-1400).

**Live cell imaging**

To examine the temporal dynamics of cellular processes during asymmetric GSC divisions, we conducted live cell imaging with high temporal resolution (e.g. 30sec, 60sec or 5min interval as mention in the figure legend and supplemental movie legend). To perform live cell imaging, adult *Drosophila* testes were dissected in a medium containing Schneider's insect medium with

200 µg/ml insulin, 15% (vol/vol) FBS, 0.6x pen/strep, with pH value at approximately 7.0, which we called “live cell medium”. Testes were then placed on a Poly-D-lysine coated FluoroDish (World Precision Instrument, Inc.), which contains the live cell medium as described. All movies were taken using spinning disc confocal microscope (Zeiss) equipped with an evolve™ camera (Photometrics), using a 63x Zeiss objective (1.4 NA). The ZEN 2 software (Zeiss) was used for acquisition with 2x2 binning. Mitotic cells were used to reconstruct 3-D movies using Imaris software (Bitplane). All videos for live cells are shown in Movie S1 – Movie S7 and Movie S10 - Movie S11.

**The 3D quantification for both time-lapse movies and fix images**

To quantify the amount of proteins segregated during asymmetric GSC divisions and symmetric SG divisions, we conducted a 3D quantification by measuring the fluorescence signal in each plane from the Z-stack (e.g. Figure S1A). The 3D quantification was done at different cell cycle stages (as labeled in the corresponding figures) in GSCs and SGs (e.g. 8-cell cyst), using time lapse movie with *cid-Dendra2*, *Dendra2-cal1*, *Dendra2-Ndc80* knock-in lines, *CID-GFP*, *α-tubulin-GFP*, and *UASp-FRT-H3-mCherry -PolyA-FRT-H3-eGFP* transgenic lines, respectively. No antibody was added to enhance Dendra2, GFP or mCherry signals for quantification.

The fixed immunostaining images used fluorescence signal of Dendra2 (CID, CAL1, and NDC80) or antibodies recognizing endogenous CID.

The 3D quantification of the fluorescence signal was done manually. Un-deconvolved raw images as 2D Z-stacks were saved as un-scaled 16-bit TIF images, and the sum of the gray values of pixels in the image (“RawIntDen”) was determined using Fiji (Image J). A circle was drawn to include all fluorescence signal (marked by Dendra2, GFP or mCherry), and an identical circle was drawn in the hub region as the background. The gray values of the fluorescence signal pixels for each Z-stack (Foreground signal, Fs) was calculated by subtracting the gray values of the background signal pixels (Background signal, Bs) from the gray values of the raw signal pixels (Raw signal, Rs). The total amount of the fluorescence signal in the nuclei was calculated by adding the gray values of the fluorescence signal from all Z-stacks. The total amount of the fluorescence signal (Fs) in the nuclei with Z-stacks (Z1+…+Zn): Fs (Z1+…+Zn) = [(Rs-Bs)1+…+(Rs-Bs)n].

**Define different categories of symmetric *versus* asymmetric CID and NDC80 at sister chromatids in prophase or prometaphase cells**

To define different categories (highly asymmetric, medium asymmetric, and symmetric) for sister centromeres in prometaphase, we used those ratios in symmetrically dividing SGs (SG1/SG2) to define the symmetric range. For example, in prometaphase, ratios above {mean + SE [1.161 + 0.015 = 1.176 (~1.2)]} are called 'asymmetric' and below are called 'symmetric'. Among the asymmetric group, we further classified them into two categories, medium asymmetric (between 1.2- to 1.4-fold difference) and highly asymmetric (> 1.4-fold difference).

To define different categories (highly asymmetric, medium asymmetric, and symmetric) for sister kinetochores in prometaphase, we used those ratios in symmetrically dividing SGs (SG1/SG2) to define the symmetric range. For example, in prometaphase, ratios above {mean + SE [1.19 + 0.02 = 1.21 (~ 1.2)]} are called 'asymmetric' and below are called 'symmetric'. Among the asymmetric group, we further classified them into two categories, medium asymmetric (between 1.2- to 1.4-fold difference) and highly asymmetric (> 1.4-fold difference).

**Define different categories of symmetric, conventional asymmetric and inverted asymmetric CID at sister chromatids in anaphase or telophase cells**

For CID inheritance patterns in anaphase-to-early telophase, ratios above {mean + SE [1.056 + 0.017 = 1.073 (~1.1)]} are called ‘asymmetric’ and below are called ‘symmetry.

To define different categories of symmetric, conventional asymmetric and inverted asymmetric for CID inheritance in anaphase or early telophase GSCs, we used those ratios in symmetrically dividing SGs (SG1/SG2) to define the symmetric range. For example, in anaphase or early telophase, ratios above {mean + SE [1.056 + 0.017 = 1.073 (~1.1)]} are called ‘conventional asymmetry’, ratios below {SG2/SG1 mean - SE [0.952 - 0.017 = 0.935 (~0.9)]} are called ‘inverted asymmetry’, and ratio between 0.9 – 1.1 are called ‘symmetry’.

**Knockdown of *cal1* (*cal1* KD) in the adult flies**

The *UAS-cal1-shRNA* flies were crossed with *nanos-Gal4 (without VP16); tub-Gal80$^{ts}$* flies to generate both *tub-Gal80$^{ts}$, nos-Gal4* (*Ctrl*) and *tub-Gal80$^{ts}$, nos-Gal4>cal1 shRNA* (*cal1* KD)] male flies. Crosses were maintained at 18°C (permissive temperature for Gal80$^{ts}$) to keep the Gal4 repressed therefore no CAL1 knockdown during development. After eclosion flies were shifted to the 29°C (restrictive temperature for Gal80$^{ts}$) for 2-10 days.

**Nocodazole Treatment Experiments**

Prior to dissection, Nocodazole (NZ) solution was freshly prepared by adding 1 µl of 2 mg/ml stock solution of NZ in DMSO per 200 µl of “live cell media” for a final NZ concentration at 10µg/ml. This solution was left in the dark at room temperature (RT) until needed. Testes were dissected in Schneider's insect media as quickly as possible and transferred to tubes where the excess media was carefully removed. After removing the Schneider's media, 50 µl of NZ solution was added to each tube, which was left open in darkness at RT. Every hour, the old NZ solution was removed and 50 µl of new NZ solution was added for a total of three exchanges or four total hours in NZ solution. At the end of the four hours, the NZ solution was removed and 1 ml of Schneider's insect media was added to each tube except the batch for immediate fixation after NZ, for which 4% formaldehyde was added to the tube instead. For the tubes where NZ was washed out, the media was removed and fresh media was added, repetitively for a total of at least five washes within 15 minutes. Testes were then fixed immediately after washout to catch prophase cells (15 min after release), prometaphase or metaphase cells (30 min after release), metaphase or anaphase cells (45 min after release), and anaphase or telophase cells (60 to 75 min after release), see Figures 7A and S7A.

**Nocodazole Recovery Experiments**

To test how cells could recover upon washing out NZ, testes were incubated in live cell media with NZ (as described above) for 2 hrs, NZ was then washed out using live cell medium (without NZ), and then incubated in live cell medium (without NZ) for at least 5hrs before imaging. Time lapse movie shows asymmetric microtubule activity could recover in GSCs (Figure 7D) and asymmetric old histone H3 segregation could partially recover (Figure S7M).

**Heat shock scheme**

Flies with *UASp*-dual color histone transgenes were paired with the *nos-Gal4* driver. Flies were raised at 18°C throughout development until adulthood to avoid pre-flip (Tran et al., 2012).

Before heat shock, 1-3 day old males were transferred to vials that had been air dried for 24 hours. Vials were submerged underneath water up to the plug in a circulating 37°C water bath for 90 minutes and recovered in a 29°C incubator for indicated time before dissection for immunostaining experiments.

**Supplementary figures and figure legends:**

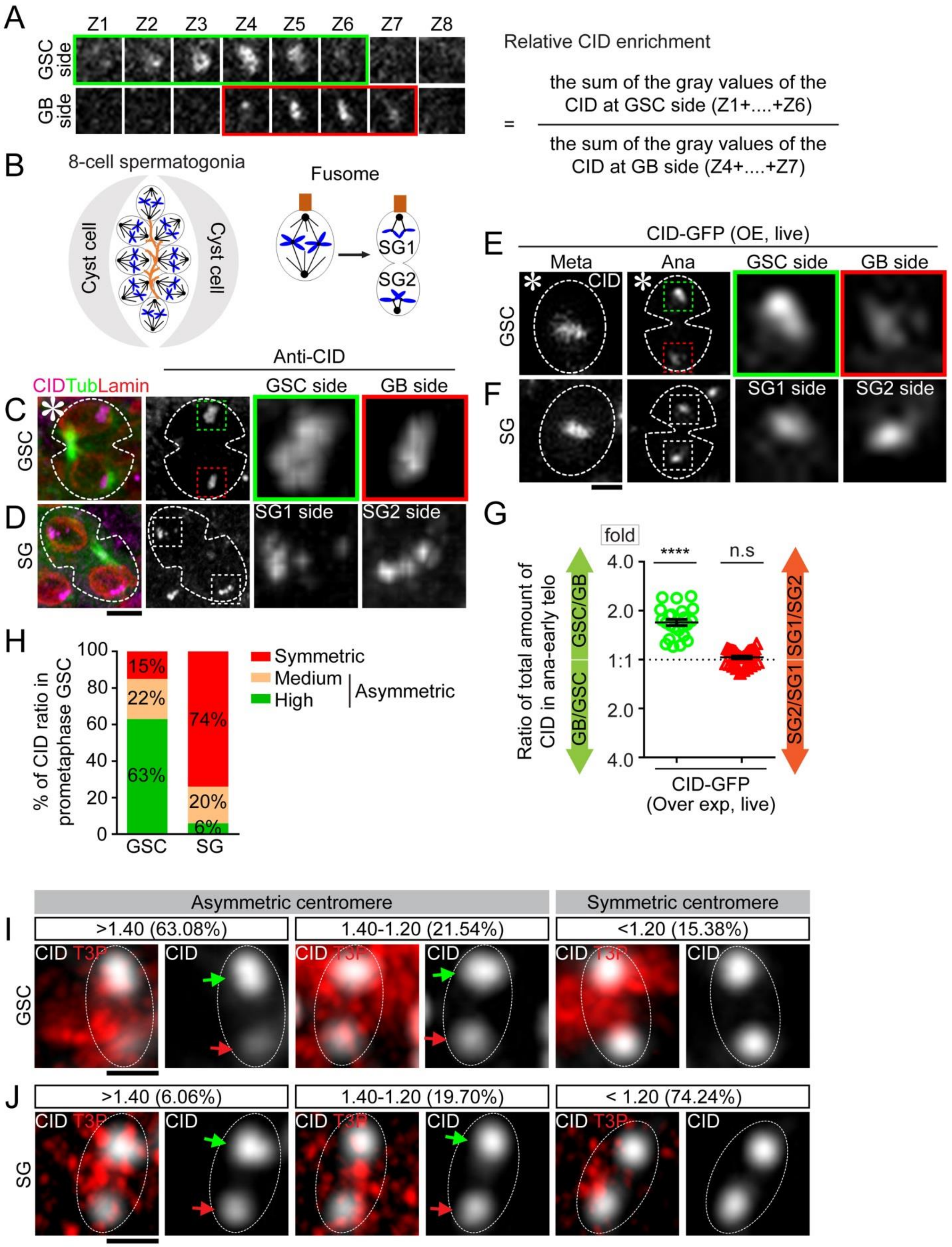

A
Z1 Z2 Z3 Z4 Z5 Z6 Z7 Z8
GSC side
GB side
Relative CID enrichment
= the sum of the gray values of the CID at GSC side (Z1+....+Z6) / the sum of the gray values of the CID at GB side (Z4+....+Z7)
B
8-cell spermatogonia
Cyst cell
Cyst cell
Fusome
SG1
SG2
C
D
CIDTubLamin
Anti-CID
GSC side
GB side
GSC
SG1 side
SG2 side
SG
E
F
CID-GFP (OE, live)
Meta
Ana
GSC side
GB side
CID
SG1 side
SG2 side
G
fold
****
n.s
Ratio of total amount of CID in ana-early telo
GSC/GB
GB/GSC
SG1/SG2
SG2/SG1
4.0
2.0
1:1
2.0
4.0
CID-GFP
(Over exp, live)
H
% of CID ratio in prometaphase GSC
100
80
60
40
20
0
15%
22%
63%
74%
20%
6%
GSC
SG
Symmetric
Medium
High
Asymmetric
I
Asymmetric centromere
Symmetric centromere
>1.40 (63.08%)
1.40-1.20 (21.54%)
<1.20 (15.38%)
CID T3P
CID
J
>1.40 (6.06%)
1.40-1.20 (19.70%)
< 1.20 (74.24%)

**Figure S1: Quantification of CID inheritance pattern in mitotic male germ cells.** (**A**) Illustration of 3D quantification: total amount of CID toward GSC side or GB side was obtained by summing CID signals from all slices with signals from a Z-stack, with background subtracted from each slice, for example, Z1 through Z6 from the GSC side and Z4 through Z7 from the GB side. The ratio was subsequently deducted by dividing the total amount of CID from GSC side by the total amount of CID from GB side. (**B**) A cartoon depicting symmetric spermatogonia cell (SG) division. Here SG1 is defined as the one in proximity to the fusome structure while SG2 is the one distal to the fusome. (**C-D**) Images of an early telophase GSC (**C**) and a SG at the same stage (**D**) from *nanos-Gal4*; *UAS-α-tubulin-GFP* testes, immunostained with anti-CID (magenta) and anti-Lamin B (red). (**E-F**) Snapshots from live cell imaging using a *cid-GFP* line, both metaphase and early anaphase were shown for a GSC (**E**) and a SG (**F**). Enlarged images show CID-GFP signals in the anaphase GSC in (**E**) and the anaphase SG in (**F**). (**G**) Quantification of CID-GFP at anaphase or early telophase GSCs and SGs, using method shown in (**A**1.73± 0.08-fold for GSC/GB (*n*= 23), 1.02± 0.02-fold for SG1/SG2 (*n*= 31), Table S3. (**H**) Percentage of different categories of symmetric (< 1.20-fold), medium asymmetric (1.20- to 1.40-fold) and highly asymmetric sister centromeres (> 1.40-fold) for GSCs (*n*= 65) and SGs (*n*= 66) in prometaphase, examples of resolved individual sister centromeres are shown in (**I**) GSCs and (**J**) SGs. Ratio= Avg± SE; *P*-value: paired *t* test. ****: $P < 10^{-4}$; n.s: no significant difference from 1:1 ratio. Asterisk: hub. Scale bars: 2μm.

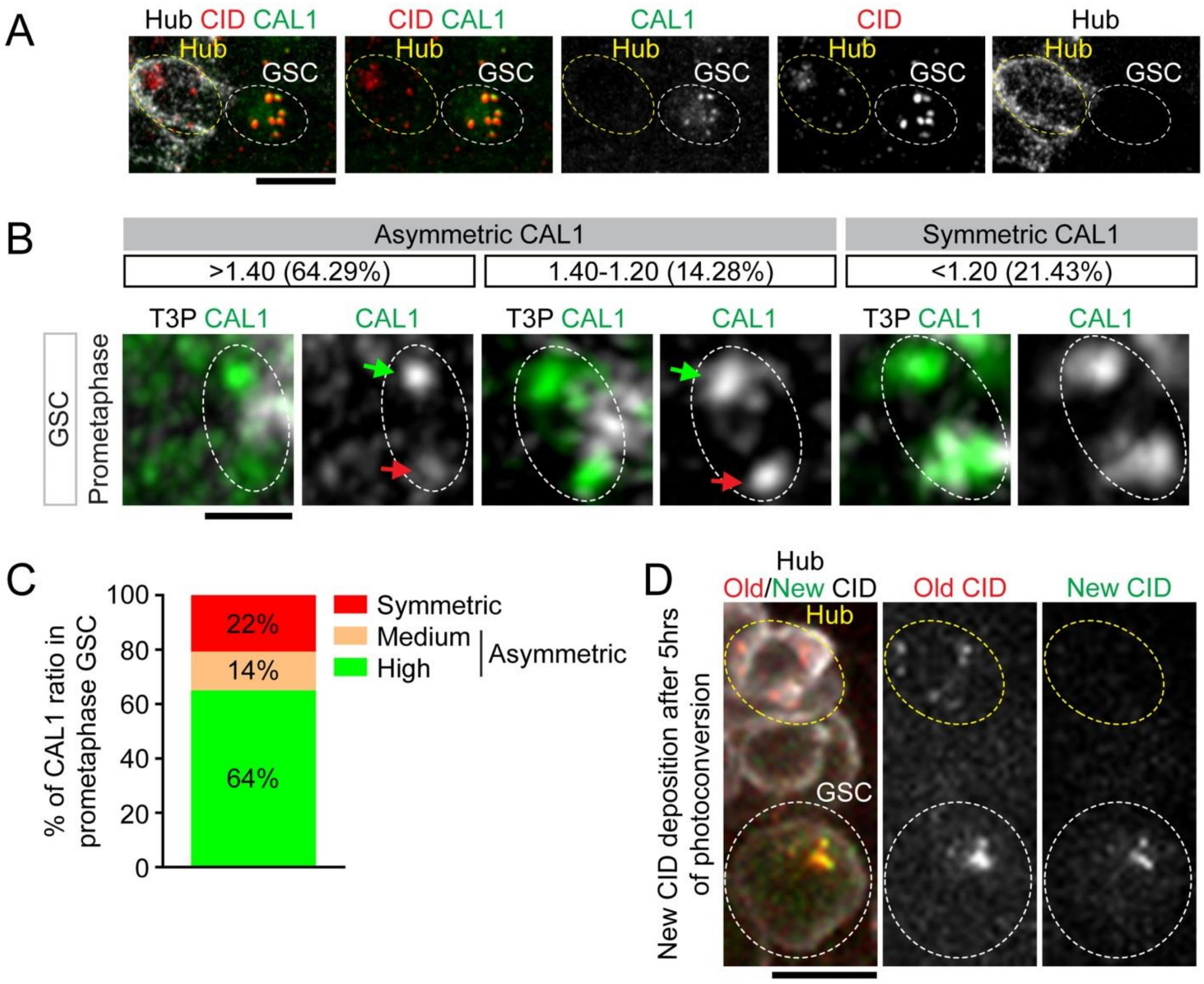

**Figure S2: CAL1 expression pattern and asymmetric CAL1 at sister centromeres in male GSCs.** (**A**) An example of the apical tip of testis from *Dendra2-cal1* knock-in male fly: Dendra2-CAL1 (green) was detectable at centromeres colocalized with CID (red) in GSCs, but was not detectable in non-replicative hub cells even though CID was detectable in hub cells, indicating specificity of CAL1 in replicative cells likely for deposition of new CID in preparation for cell division. (**B**) In prometaphase GSCs, distribution patterns of CAL1 at resolved individual sister centromeres showed different categories of asymmetry: highly asymmetric (> 1.40-fold), medium (between 1.2-1.4-fold) and (<1.2-fold) in GSCs. (**C**) Quantification of percentages of different categories in GSCs (*n*= 28). (**D**) After photoconversion, testes were cultured for 5hrs before fixation. New CID (green Dendra2) incorporation is detectable in early prophase GSCs but not in non-replicative hub cells. Scale bars: 5μm (**A, D**) and 0.5μm (**B**).

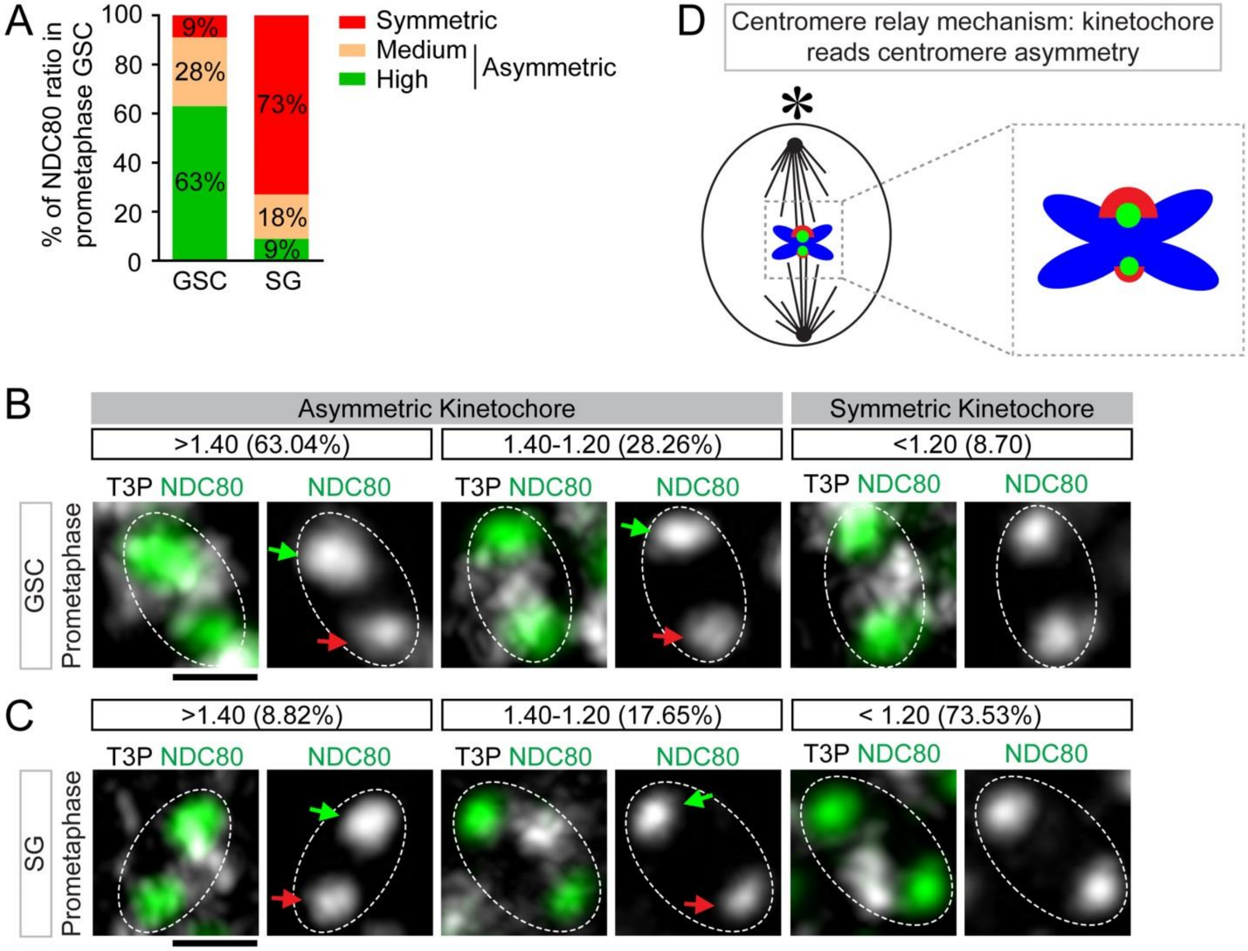

**Figure S3: Distribution of a key kinetochore protein NDC80 at sister kinetochores.** (**A**) Percentage of different categories of symmetric (< 1.20-fold), medium asymmetric (1.20- to 1.40-fold) and highly asymmetric sister kinetochores (> 1.40-fold) for GSCs (*n*= 46) and SG (*n*= 34) in prometaphase, examples of resolved individual sister kinetochores are shown in (**B**) GSCs and (**C**) SGs. (**D**) A cartoon depicting a potential 'relay' mechanism: The sister centromere asymmetry is read out as the asymmetry between sister kinetochores. Asterisk: hub. Scale bars: 0.5μm.

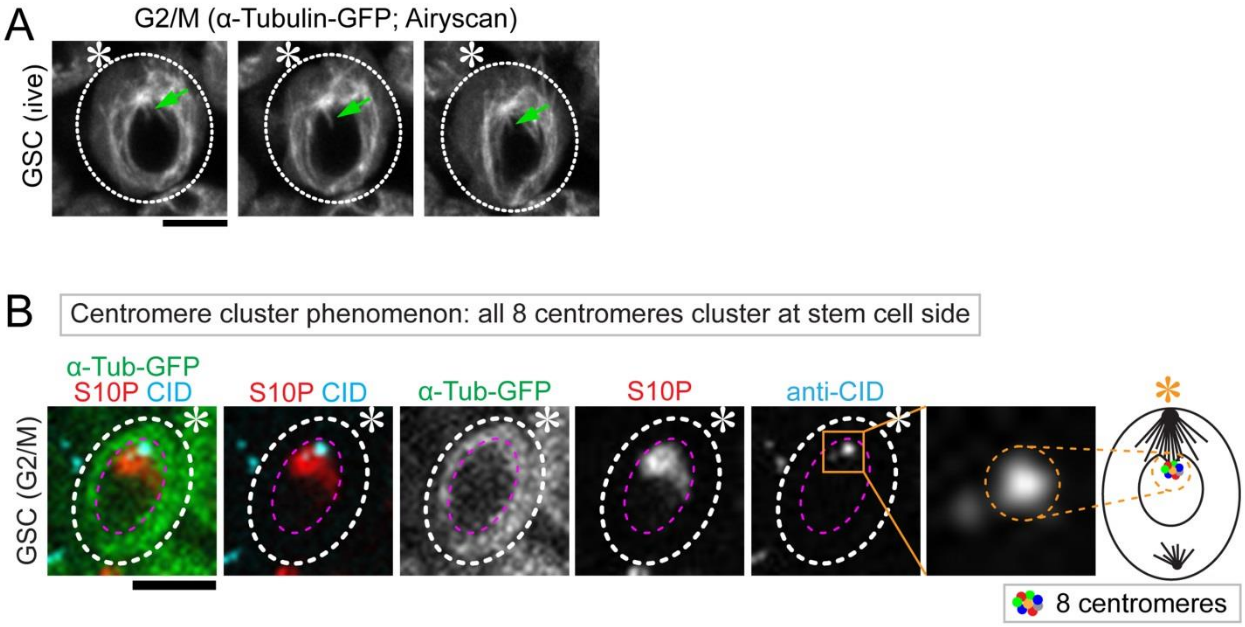

**Figure S4: Intimate interactions between microtubules and centromeres in GSCs.** (**A**) The ‘poking in’ activities of microtubules from high resolution live cell imaging (Airyscan) using the *nanos-Gal4*; *UAS-α-tubulin-GFP* line (snapshots from Movie S7). (**B**) All CID signals (blue, centromeres from all chromosomes) were clustered near the nuclear envelope toward the niche side in GSCs at G2-to-M phase transition or prophase, with detectable H3S10P (red) and α-Tubulin signal (green). Asterisk: hub. Scale bars: 5μm.

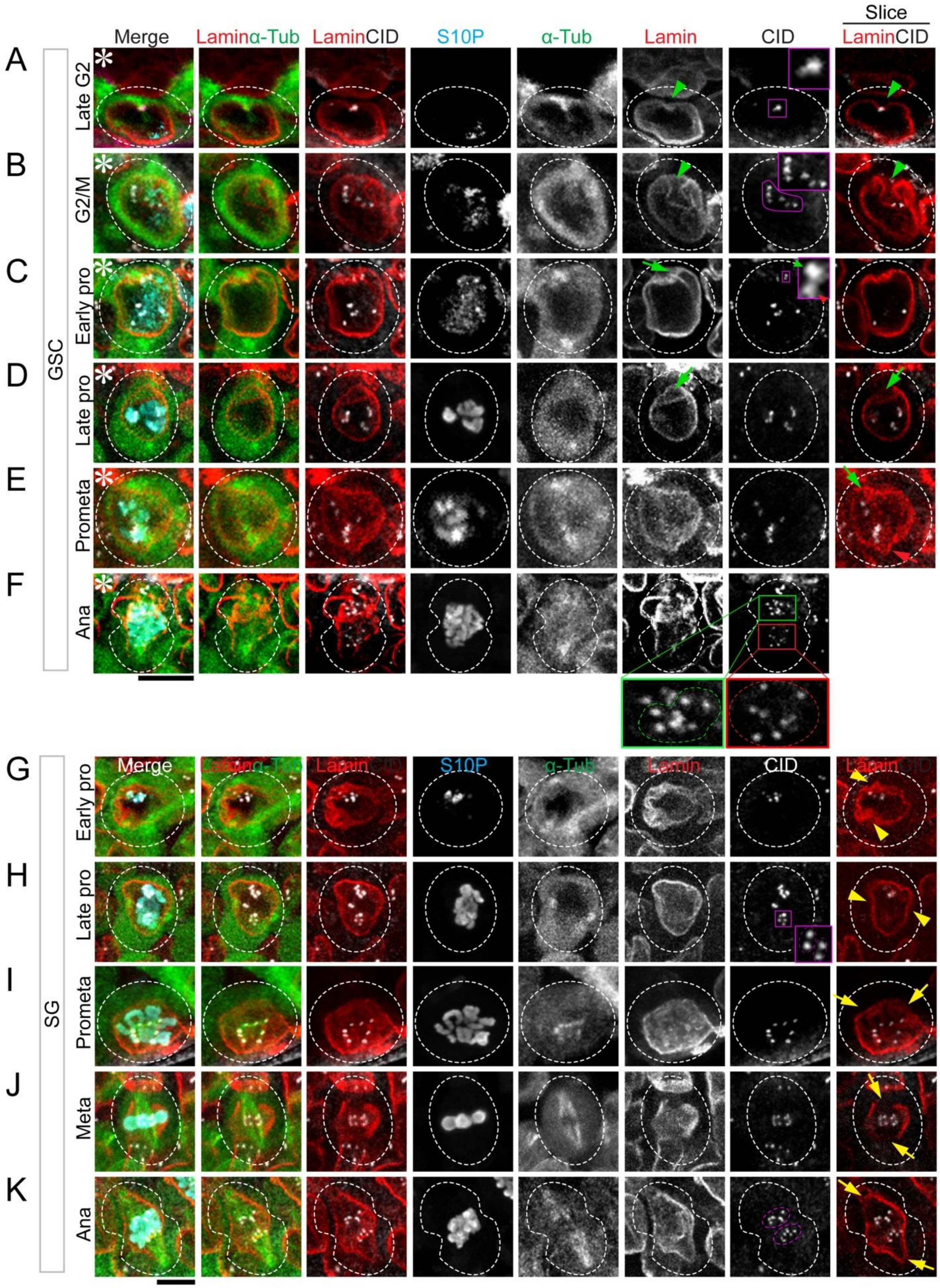

Slice
Merge
Laminα-Tub
LaminCID
S10P
α-Tub
Lamin
CID
LaminCID
A
B
C
D
E
F
G
H
I
J
K
Late G2
G2/M
Early pro
Late pro
Prometa
Ana
Meta
GSC
SG

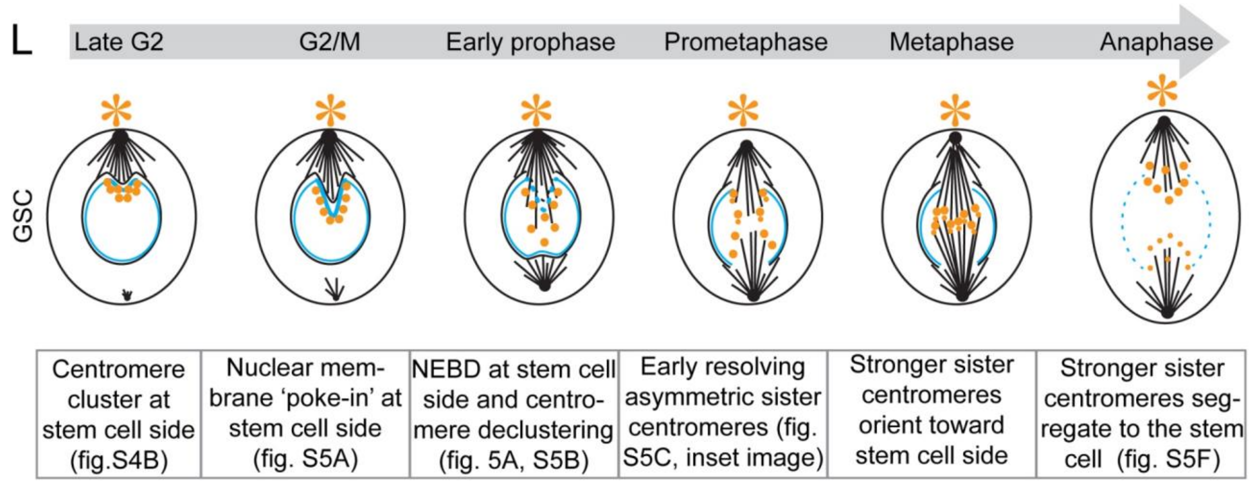

**Figure S5: Polarized NEBD potentially allows for differential attachment of microtubules to sister centromeres.** (**A-F**) Morphology of nuclear lamina at different cell cycle stages of GSCs, visualized by immunostaining with anti-Lamin B (red), co-stained with anti-CID (white) and anti-H3S10P (blue) in *nanos-Gal4*; *UAS-α-tubulin-GFP* (green) line. Polarized nuclear lamina invagination and centromere cluster at GSC side was labeled by green arrowheads from late G2 to G2/M transition (**A-B**). The ‘poking in’ activities of microtubules lead to centromere declustering along nuclear membrane (**B**). The ‘poking in’ activities of microtubules labeled by green arrow (from GSC side) at prophase to prometaphase and red arrow (from GB side) at prometaphase (**C-E**). Inset at early prophase shows a pair of resolved asymmetric sister centromeres (**C**). (**F**) Inset at anaphase show stronger sister centromeres segregated to the GSC side (green outlines), compared to the sister centromeres segregated to the GB side (red outlines). (**G-K**) Morphology of nuclear lamina at different cell cycle stages of SGs, visualized by immunostaining with anti-Lamin B (red), co-stained with anti-CID (white) and anti-H3S10P (blue) with the *nanos-Gal4*; *UAS-α-tubulin-GFP* (green) line. Nuclear lamina invagination was detected at both sides in SG labeled by yellow arrowheads at prophase (**G-H**). Inset at late prophase show two pairs of resolved symmetric sister centromeres (**H**). The ‘poking in’ activities of microtubules were from both poles labeled by yellow arrows at prometaphase and metaphase (**I-J**). At anaphase, segregated sister centromeres show a symmetric pattern in SG (**K**). **(L)** A cartoon depicting the sequential events of the 'mitotic drive' in GSCs. Asterisk: hub. Scale bars: 5μm.

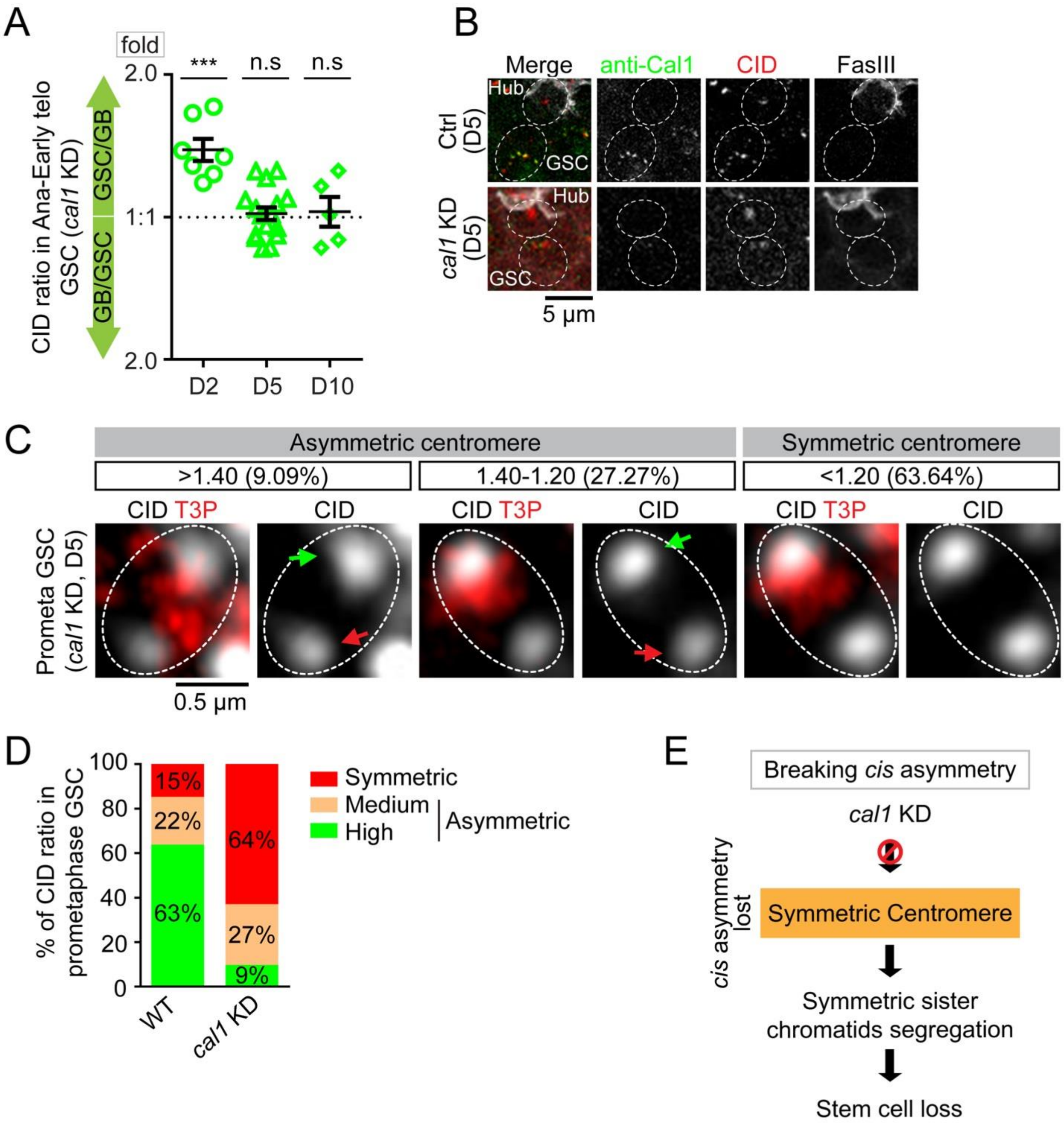

**Figure S6: Breaking *cis*-asymmetry of CID at sister centromeres by knocking down CAL1.** **(A)** Gradual loss of asymmetric CID inheritance patterns in anaphase to early telophase GSCs, from *cal1* KD testes two days (D2), five days (D5) or ten days (D10) at the restrictive temperature (29°C). (**B**) The efficiency of *cal1* KD was quantified at D5 and D10 using immunostaining with anti-CAL1. (**C**) Examples of resolved individual sister centromeres are shown in *cal1* KD GSCs. (**D**) Percentage of different categories of symmetric (< 1.20-fold), medium asymmetric (1.20- to 1.40-fold) and highly asymmetric sister centromeres (> 1.40-fold) for wild-type (WT) GSCs (*n*= 65) and *cal1* KD GSCs (D5, *n*=33) in prometaphase. **(E)** A cartoon shows breaking *cis* asymmetry by compromising CAL1 and the consequences of symmetric sister centromere establishment and segregation, as well as stem cell loss.

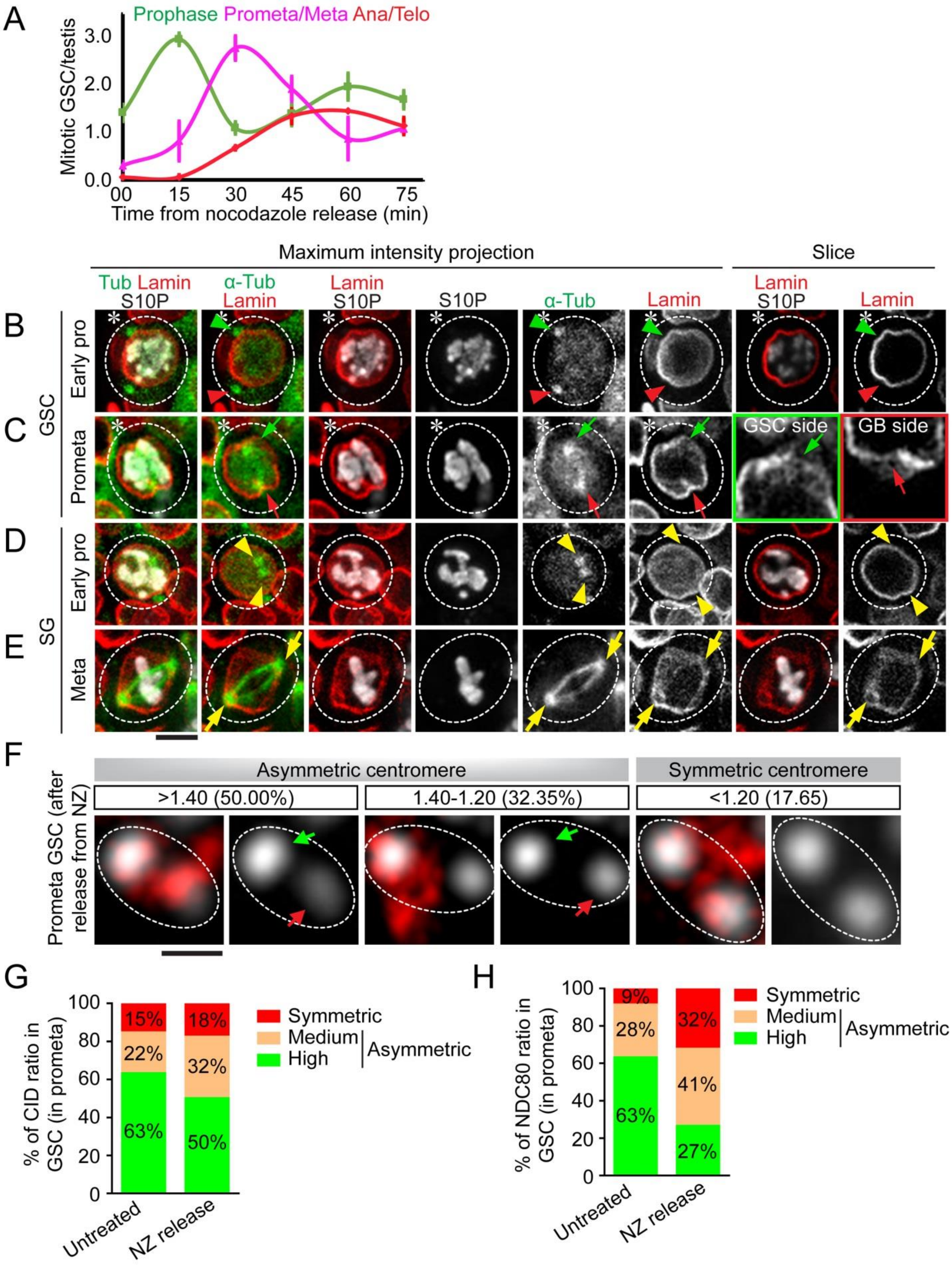
A
Prophase Prometa/Meta Ana/Telo
Mitotic GSC/testis
3.0
2.0
1.0
0.0
00 15 30 45 60 75
Time from nocodazole release (min)
Maximum intensity projection
Slice
Tub Lamin S10P
α-Tub Lamin
Lamin S10P
S10P
α-Tub
Lamin
Lamin S10P
Lamin
B
C
D
E
GSC
SG
Early pro
Prometa
Early pro
Meta
GSC side
GB side
F
Asymmetric centromere
Symmetric centromere
>1.40 (50.00%)
1.40-1.20 (32.35%)
<1.20 (17.65)
Prometa GSC (after release from NZ)
G
% of CID ratio in GSC (in prometa)
15% 22% 63%
18% 32% 50%
Untreated
NZ release
Symmetric
Medium
High
Asymmetric
H
% of NDC80 ratio in GSC (in prometa)
9% 28% 63%
32% 41% 27%
Untreated
NZ release
Symmetric
Medium
High
Asymmetric

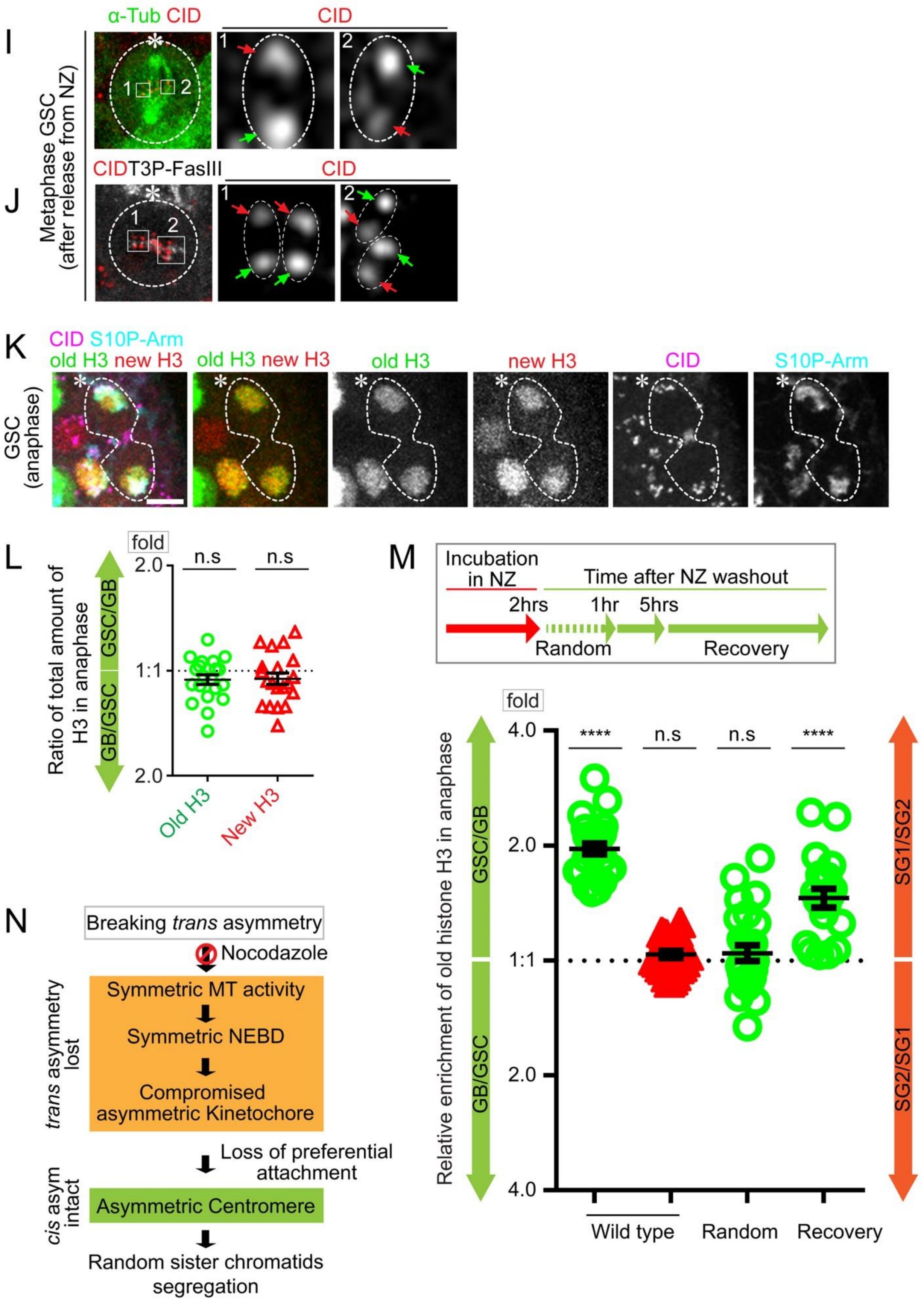

I
α-Tub CID
CID
Metaphase GSC (after release from NZ)
J
CID T3P-FasIII
CID
K
CID S10P-Arm
old H3 new H3
old H3 new H3
old H3
new H3
CID
S10P-Arm
GSC (anaphase)
L
fold
n.s
n.s
2.0
1:1
2.0
GSC/GB
GB/GSC
Ratio of total amount of H3 in anaphase
Old H3
New H3
M
Incubation in NZ
Time after NZ washout
2hrs
1hr
5hrs
Random
Recovery
fold
4.0
2.0
1:1
2.0
4.0
****
n.s
n.s
****
GSC/GB
GB/GSC
SG1/SG2
SG2/SG1
Relative enrichment of old histone H3 in anaphase
Wild type
Random
Recovery
N
Breaking *trans* asymmetry
Nocodazole
Symmetric MT activity
Symmetric NEBD
Compromised asymmetric Kinetochore
*trans* asymmetry lost
Loss of preferential attachment
*cis* asym intact
Asymmetric Centromere
Random sister chromatids segregation

**Figure S7: Breaking *trans*-asymmetry of microtubules using nocodazole (NZ) treatment.** (**A**) GSCs are primarily arrested at G2/M or early prophase with NZ treatment. Then immediately after washing out NZ, GSCs progress to prophase after 15 minutes, prometaphase to metaphase after 30 minutes, and anaphase to telophase after 45 to 60 minutes. All time points = Avg$\pm$ SE (*n*= 28 testes for t= 0 min, *n*= 22 testes for t= 15 min, *n*= 22 testes for t= 30 min, *n*= 24 testes for t= 45 min, *n*= 22 testes for t= 60 min, and *n*= 24 testes for t= 75 min). (**B-E**) Morphology of nuclear envelope visualized by immunostaining using anti-Lamin B (red), co-stained with anti-H3S10P (white) using *nanos-Gal4*; *UAS-$\alpha$-tubulin-GFP* line (green): in GSCs at early prophase (**B**) and at prometaphase (**C**), when symmetric microtubule activity from both centrosomes could be visualized by $\alpha$-Tubulin-GFP in GSCs after releasing from NZ-induced cell cycle arrest; in SGs at early prophase (**D**) and at metaphase (**E**). (**F**) In prometaphase GSCs immediately after releasing from NZ arrest (*n*= 34), examples of sister centromeres with different categories: symmetric (< 1.20-fold), medium asymmetric (1.20- to 1.40-fold) and highly asymmetric (> 1.40-fold) resolved individual sister centromeres. (**G**) Percentages of CID ratios at resolved individual sister centromeres in prometaphase with different categories in untreated GSCs (*n*= 65) and GSCs immediately after releasing from NZ arrest (*n*= 34). (**H**) Percentages of NDC80 ratios at resolved individual sister centromeres in prometaphase with different categories in untreated GSCs (*n*= 65) and GSCs immediately after releasing from NZ arrest (*n*= 34). (**I-J**) Immediately after releasing from NZ arrest, sister chromatids with asymmetric sister centromeres are randomly attached by microtubules from mother centrosome *versus* daughter centrosome at the metaphase plate in two representative GSCs (**I** and **J**): examples include some stronger centromeres are attached to the microtubule emanating from the daughter centrosome (pattern 1 in **I-J**, green arrows) and some stronger centromeres are attached to the microtubule emanating from the mother centrosome (pattern 2 in **I-J**, green arrows). (**K-L**) Random histone H3 inheritance patterns in anaphase GSCs immediately after releasing from NZ-induced cell cycle arrest (**K**), as quantified in (**L**): old H3-GFP GSC/GB= 0.95$\pm$ 0.03; new H3-mKO GB/GSC= 0.96$\pm$ 0.04, ($n$= 20), Table S17. (**M**) Quantification of old histone H3 inheritance pattern in anaphase cells from live cell imaging: wild-type GSCs (old H3-mCherry GSC/GB= 1.98$\pm$ 0.07, $n$= 26), wild-type SGs (old H3-mCherry SG1/SG2= 1.04$\pm$ 0.02, $n$= 25), GSCs immediately after releasing from NZ arrest show random pattern (old H3-mCherry GSC/GB= 1.08$\pm$ 0.06, $n$= 26), and some GSC recovered from NZ arrest show asymmetry again (old H3-mCherry GSC/GB= 1.50$\pm$ 0.09, $n$= 20), Table S18. (**N**) A cartoon shows breaking *trans* asymmetry by depolymerizing microtubules and the consequences of loss of preferential microtubule attachment as well as random sister chromatids segregation. Ratio= Avg$\pm$ SE; $P$-value: paired $t$ test. ****: $P< 10^{-4}$; n.s: no significant difference. Asterisk: hub. Scale bars: 5μm.

**Supplementary movie legends:**

**Movie S1: Asymmetric CID-Dendra2 segregation in GSC.** Live cell imaging of the CID-Dendra2 in a *Drosophila* male GSC: This video is a 3D reconstruction of individual images (8 × 1-μm interval optical sections per frame), showing asymmetric CID-Dendra2 segregation during asymmetric GSC division. The video was acquired at 30sec intervals for 5-6 minutes from prophase to anaphase. Metaphase is used as a landmark to define time point zero, and other time points are labeled as minus minutes prior to metaphase. The quantification is shown in Figure 1D. Asterisk: hub. Scale bar: 2μm.

**Movie S2: Symmetric CID-Dendra2 segregation in SG.** Live cell imaging of the CID-Dendra2 in a *Drosophila* male SG: This video is a 3D reconstruction of individual images (8 × 1-μm interval optical sections per frame), showing symmetric CID-Dendra2 segregation during symmetric SG division. The video was acquired at 30sec intervals for 5-6 minutes from prophase to anaphase. Metaphase is used as a landmark to define time point zero, and other time points are labeled as minus minutes prior to metaphase. The quantification is shown in Figure 1D. Scale bar: 2μm.

**Movie S3: Asymmetric CID-GFP segregation in GSC.** Live cell imaging of CID-GFP in a *Drosophila* male GSC: This video is a 3D reconstruction of individual images (8 × 1-μm interval optical sections per frame), showing asymmetric CID-GFP segregation during asymmetric GSC division. The video was acquired at 30sec intervals for 5-6 minutes from prophase to anaphase. Metaphase is used as a landmark to define time point zero, and other time points are labeled as minus minutes prior to metaphase. The quantification is shown in Figure S1G. Asterisk: hub. Scale bar: 5μm.

**Movie S4: Symmetric CID-GFP segregation in SG.** Live imaging of the CID-GFP in a *Drosophila* male SG: This video is a 3D reconstruction of individual images (8 × 1-μm interval optical sections per frame), showing symmetric CID-GFP segregation during symmetric SG division. The video was acquired at 30sec intervals for 5-6 minutes from prophase to anaphase. Metaphase is used as a landmark to define time point zero, and other time points are labeled as minus minutes prior to metaphase. The quantification is shown in Figure S1G. Scale bar: 5μm.

**Movie S5: Temporal asymmetry of α-Tubulin in GSC**. Live cell imaging of α-Tubulin-GFP in a *Drosophila* male GSC: This video is a 3D reconstruction of individual images (14 × 1-μm interval optical sections per frame), showing temporal asymmetry in α-Tubulin-GFP activity at mother centrosome (labeled by green arrow) and daughter centrosome (labeled by red arrow). The mother centrosome is actively nucleating microtubules from mid- to late G2 phase (-240min to -50min), whereas the daughter centrosome nucleates microtubules upon mitotic entry (~ -50 min). Also, in prophase, the dominance of mother centrosome activity becomes less, while microtubules from the daughter centrosome start to increase (-35min to -5min). However, at metaphase microtubules from both sides become equal (0 min). The video was acquired at 5-min interval for 5-6 hrs from mid G2 to G1. Metaphase is used as a landmark to define time point zero, and other time points are labeled as minus minutes prior to metaphase. The quantification is shown in Figure 3C. Asterisk: hub. Scale bar: 5μm.

**Movie S6: Symmetric α-tubulin in SG**. Live cell imaging of the α-tubulin-GFP in a *Drosophila* male SG: This video is a 3D reconstruction of individual images (8 × 1-μm interval optical sections per frame), showing neither the mother nor the daughter centrosome nucleates microtubules throughout G2 phase. However, both centrosomes start nucleating microtubules simultaneously upon mitotic entry (~ -35min). The video was acquired at 5-min interval for 5-6 minutes from G2/M to G1. Metaphase is used as a landmark to define time point zero, and other time points are labeled as minus minutes prior to metaphase. The quantification is shown in Figure 3D. Scale bar: 5μm.

**Movie S7: A high-resolution time lapse movie of GSC to visualize asymmetric microtubule 'poking in' phenomenon**. Live cell imaging of the α-tubulin-GFP in a *Drosophila* male GSC using LSM800 confocal microscopy equipped with Airyscan: This video is a 3D reconstruction of individual images (6 × 0.15μm interval optical sections per frame), showing microtubules spike "poking in" (labeled by green arrow) at GSC side at G2/M transition. The video was acquired at 5-min interval for 30 minutes from G2/M transition to prophase. The G2/M is time point zero; snapshots are shown in Figure S4A. Asterisk: hub. Scale bar: 5μm.

**Movie S8: The 3D reconstructed GSC in early-prophase showing NEBD at the GSC side.** This video is a 3D reconstruction of individual fixed images (5 × 0.15μm interval optical sections per frame) of α-tubulin-GFP-expressing *Drosophila* male GSC at early prophase, immunostained with anti-Lamin B and anti-CID. (**A**) A 3D reconstruction with α-tubulin-GFP (green), anti-Lamin B (magenta) and anti-CID (red) signals. (**B**) A 3D reconstruction with anti-Lamin B (white) and anti-CID (red) signals. (**C**) A zoom-in view of (**B**). This 3D reconstruction shows asymmetric nuclear envelope breakdown at the GSC side in early prophase. The maximum projection of the image is shown in Figure 4A. Asterisk: hub. Scale bar: 5μm.

**Movie S9: The 3D reconstructed GSC in late-prophase showing NEBD at the GSC side and "poking in" phenomenon at the GB side.** This video is a 3D reconstruction of individual fixed images (10 × 0.15μm interval optical sections per frame) of α-tubulin-GFP-expressing

*Drosophila* male GSC at late-prophase, immunostained with anti-Lamin B and anti-CID. (**A**) A 3D reconstruction with α-tubulin-GFP (green), anti-Lamin B (magenta) and anti-CID (red) signals. (**B**) A 3D reconstruction with anti-Lamin B (white) and anti-CID (red) signals. This image shows that the nuclear envelope at stem cell side is broken, and the nuclear envelope at GB side is "poking in" at late-prophase. The maximum projection of the image is shown in Figure 4B. Asterisk: hub. Scale bar: 5μm.

**Movie S10: Symmetric α-Tubulin nucleation in GSC after release from nocodazole (NZ) arrest**. Live cell imaging of α-Tubulin-GFP in a *Drosophila* male GSC released from NZ arrest: This video is a reconstruction of individual images (7 × 1-μm interval optical sections per frame), showing symmetry in α-Tubulin-GFP activity at mother centrosome (labeled by green arrow) and daughter centrosome (labeled by red arrow). Both mother and daughter centrosomes symmetrically active after release from NZ arrest, and immediately enter into prophase. The video was acquired at 5-min interval for 15-30 min after release from NZ. Metaphase is used as a landmark to define time point zero, and other time points are labeled as minus minutes prior to metaphase. The quantification is shown in Figures 7B-C. Asterisk: hub. Scale bar: 5μm.

**Movie S11: Recovery of temporal asymmetry of α-Tubulin in GSC after release from nocodazole (NZ) arrest**. Live cell imaging of α-Tubulin-GFP in a *Drosophila* male GSC: This video is a reconstruction of individual images (15 × 1-μm interval optical sections per frame), showing temporal asymmetry in α-Tubulin-GFP activity at mother centrosome (labeled by green arrow) and daughter centrosome (labeled by red arrow) is recovered ~10hrs after released from NZ arrest. The mother centrosome is actively nucleating microtubules from mid- to late G2 phase (-250min to -50min), whereas the daughter centrosome nucleates microtubules in G2/M phase (~ -50 min). Also, in prophase, the dominance of mother centrosome activity becomes less, while microtubules from the daughter centrosome start to increase (-45min to -5min). However, at metaphase microtubules from both sides become equal (0 min). The video was acquired at 5-min interval for ~10 hrs. Metaphase is used as a landmark to define time point zero, and other time points are labeled as minus minutes prior to metaphase. The montage is shown in Figure 7D. Asterisk: hub. Scale bar: 5μm.

**Supplementary tables and table legends:**

**Table S1: Quantification of CID in anaphase to early telophase GSCs and SGs by immunostaining using anti-CID.**

| # | GSC/GB | SG1/SG2 |
|---|---|---|
| **1** | 1.181852587 | 1.072688683 |
| **2** | 1.433974059 | 1.123507204 |
| **3** | 1.755702557 | 1.142589544 |
| **4** | 1.28693445 | 1.061786461 |
| **5** | 1.735516044 | 1.098328681 |
| **6** | 1.262904709 | 1.010135462 |
| **7** | 1.259372589 | 1.149435865 |
| **8** | 1.316137585 | 1.109957322 |
| **9** | 1.227829389 | 1.086620658 |
| **10** | 1.332531717 | 0.977108187 |
| **11** | 1.336723733 | 0.968406269 |
| **12** | 1.304848805 | 0.974064695 |
| **13** | 1.201435711 | 1.100481281 |
| **14** | 1.194083643 | 1.073399858 |
| **15** | 1.20879706 | 1.045534985 |
| **16** | 1.16374964 | 0.916112069 |
| **17** | 1.565665091 | 1.206757464 |
| **18** | 1.258401145 | 1.058884543 |

| | | |
|---|---|---|
| **19** | 1.191521531 | 0.925785447 |
| **20** | 1.5217 | 0.910269051 |
| **21** | 1.5701 | 1.161636493 |
| **22** | 1.587 | 1.150879872 |
| **23** | 1.531 | 0.97068742 |
| **24** | 1.686 | |
| **25** | 1.716 | |
| **26** | 1.694 | |
| **27** | 1.551 | |

**Table S2: Quantification of CID-Dendra2 in anaphase to early telophase GSCs and SGs by live cell imaging.**

| # | GSC/GB | SG1/SG2 |
|---|---|---|
| 1 | 1.351036022 | 1.142898421 |
| 2 | 1.151915643 | 1.18709256 |
| 3 | 1.635341655 | 1.155233976 |
| 4 | 1.325772996 | 1.024702142 |
| 5 | 1.243987285 | 1.022841092 |
| 6 | 1.226437723 | 1.091847321 |
| 7 | 1.522769663 | 1.046892536 |
| 8 | 1.335211721 | 1.004702142 |
| 9 | 1.333865917 | 1.002841092 |
| 10 | 1.694542979 | 1.071847321 |
| 11 | 1.616764902 | 1.026892536 |
| 12 | 1.355781874 | 0.882695936 |
| 13 | 1.453536058 | 0.960143503 |
| 14 | 1.244966152 | 0.942263279 |
| 15 | 1.268494034 | 0.924058226 |
| 16 | 1.220751075 | 0.995289367 |
| 17 | 1.413726448 | 1.108976872 |
| 18 | 1.263630877 | 1.153306665 |
| 19 | 1.531092331 | 1.036479976 |
| 20 | 1.531092331 | 1.058431 |
| 210 | 1.51757744 | 0.978612 |
| 22 | 1.695118043 | |
| 23 | 1.150776217 | |

**Table S3: Quantification of CID-GFP in anaphase to early telophase GSCs and SGs by live cell imaging.**

| # | GSC/GB | SG1/SG2 |
|---|---|---|
| 1 | 1.628534474 | 1.126563215 |
| 2 | 1.891987337 | 1.25080379 |
| 3 | 1.870089138 | 0.855378707 |
| 4 | 2.412713879 | 0.909605149 |
| 5 | 1.807891532 | 0.932446439 |
| 6 | 1.616359093 | 0.820882642 |
| 7 | 1.982243242 | 1.05901652 |
| 8 | 1.734670914 | 1.141760345 |
| 9 | 1.517357554 | 1.137522276 |
| 10 | 2.453457296 | 0.982201725 |
| 11 | 1.301026153 | 1.03806825 |
| 12 | 1.577669942 | 0.941670065 |
| 13 | 1.221849712 | 0.939137303 |
| 14 | 1.245031491 | 1.156874829 |
| 15 | 1.407052752 | 1.140599551 |
| 16 | 1.203254498 | 1.083258785 |
| 17 | 1.331363218 | 0.883599481 |
| 18 | 2.055 | 0.958347113 |
| 19 | 2.387 | 0.91445709 |
| 20 | 1.756 | 1.055 |
| 210 | 2.0287 | 1.1616 |
| 22 | 1.644 | 1.1686 |
| 23 | 1.689 | 1.038 |
| 24 | | 1.1311 |
| 25 | | 1.1204 |
| 26 | | 1.025 |
| 27 | | 1.136 |
| 28 | | 1.0961 |
| 29 | | 1.134 |
| 30 | | 0.8813 |
| 31 | | 0.9719 |

**Table S4: Quantification of sister centromeres in prometaphase GSCs and SGs by immunostaining using anti-CID.**

| Sister centromere pair # | GSC | SG |
|---|---|---|
| 1 | 1.454398955 | 1.0634278 |
| 2 | 1.581907307 | 1.3777814 |
| 3 | 2.176156234 | 1.140667 |
| 4 | 1.741236547 | 1.0315564 |
| 5 | 1.699125876 | 1.1013624 |
| 6 | 1.255762531 | 1.4031346 |
| 7 | 1.351585293 | 1.052882 |
| 8 | 1.777628993 | 1.1007956 |
| 9 | 1.704303192 | 1.1496751 |
| 10 | 1.795453231 | 1.022172 |
| 11 | 1.421553123 | 1.0104498 |
| 12 | 1.517118421 | 1.0231168 |
| 13 | 2.41321584 | 1.1063371 |
| 14 | 1.821021564 | 1.1807855 |
| 15 | 1.287924214 | 1.2682486 |
| 16 | 1.4460641 | 1.0095048 |
| 17 | 1.522308253 | 1.095408 |
| 18 | 1.703342201 | 1.2023619 |
| 19 | 1.548014856 | 1.4595929 |
| 20 | 1.808520217 | 1.149548 |
| 21 | 1.538352082 | 1.1998526 |
| 22 | 1.226093901 | 1.0630755 |
| 23 | 1.177096933 | 1.0897182 |
| 24 | 1.838649193 | 1.2367325 |
| 25 | 1.118389059 | 1.4739724 |
| 26 | 1.045663473 | 1.1731339 |
| 27 | 1.823550139 | 1.0046856 |
| 28 | 1.548014856 | 1.0534605 |
| 29 | 1.944016052 | 1.1013694 |
| 30 | 1.517747223 | 1.2635175 |
| 31 | 1.540362533 | 1.074951 |
| 32 | 1.375114159 | 1.103874 |
| 33 | 1.08538421 | 1.0813608 |
| 34 | 1.056765708 | 1.33895 |
| 35 | 1.150759142 | 1.1163995 |
| 36 | 1.912610051 | 1.0684896 |
| 37 | 1.150454735 | 1.0339541 |
| 38 | 1.992795765 | 1.2625603 |
| 39 | 1.5800927 | 1.1602625 |

| | | |
|---|---|---|
| **40** | 1.700652101 | 1.1922376 |
| **41** | 2.04745014 | 1.2252916 |
| **42** | 1.978423281 | 1.2730751 |
| **43** | 1.265846073 | 1.2643714 |
| **44** | 1.562712458 | |
| **45** | 1.182445488 | |
| **46** | 1.548014856 | |
| **47** | 1.944016052 | |
| **48** | 1.517747223 | |
| **49** | 1.219794276 | |
| **50** | 1.325526552 | |
| **51** | 1.378307066 | |
| **52** | 1.209003771 | |
| **53** | 1.384818529 | |
| **54** | 1.241964123 | |
| **55** | 1.180496206 | |
| **56** | 1.523968091 | |
| **57** | 1.221398675 | |
| **58** | 1.406147701 | |
| **59** | 1.467713081 | |

**Table S5: Quantification of CAL1 at sister centromeres in prometaphase GSCs using Dendra2-CAL1 knock-in line.**

| CAL1 at Pair of Sister Centromeres | GSC |
|---|---|
| 1 | 1.762847017 |
| 2 | 1.668916069 |
| 3 | 1.273918222 |
| 4 | 1.697133177 |
| 5 | 1.129874652 |
| 6 | 2.10906298 |
| 7 | 2.022021661 |
| 8 | 2.401664145 |
| 9 | 1.923152227 |
| 10 | 1.23669328 |
| 11 | 1.194784934 |
| 12 | 1.158666434 |
| 13 | 1.818111053 |
| 14 | 1.023058252 |
| 15 | 1.176882208 |
| 16 | 1.426731079 |
| 17 | 1.095514588 |
| 18 | 2.497839239 |
| 19 | 1.968957283 |
| 20 | 2.589546299 |
| 21 | 1.54187876 |
| 22 | 1.962859796 |
| 23 | 1.117746797 |
| 24 | 2.496050033 |
| 25 | 1.714984377 |
| 26 | 1.819881441 |
| 27 | 1.670472136 |
| 28 | 1.035405193 |

**Table S6: Quantification of NDC80 in anaphase to early telophase GSCs and SGs using the Dendra2-NDC80 knock-in line.**

| # | GSC/GB | SG1/SG2 |
|---|---|---|
| **1** | 1.20591 | 0.92579 |
| **2** | 1.22725 | 1.05888 |
| **3** | 1.3778 | 0.91027 |
| **4** | 1.5641 | 1.16164 |
| **5** | 1.70618 | 1.15088 |
| **6** | 1.19781 | 0.97069 |
| **7** | 1.3176 | 1.11056 |
| **8** | 1.53487 | 0.94527 |
| **9** | 1.46742 | 0.98711 |
| **10** | 1.8596 | 1.01336 |
| **11** | 1.29055 | 0.88362 |
| **12** | 1.743 | 1.01012 |
| **13** | 2.19932 | 1.16871 |
| **14** | 1.84774 | 1.15286 |
| **15** | 1.81431 | 1.08919 |
| **16** | 1.29751 | 1.32728 |
| **17** | 1.22645 | 1.02107 |
| **18** | 1.22381 | 1.13465 |
| **19** | 1.21363 | 1.17153 |
| **20** | | 0.78864 |
| **21** | | 0.80937 |
| **22** | | 0.97844 |
| **23** | | 0.83539 |
| **24** | | 1.16991 |
| **25** | | 0.99829 |
| **26** | | 1.13468 |
| **27** | | 0.99595 |
| **28** | | 1.02157 |

**Table S7: Quantification of sister kinetochores in prometaphase GSCs and SGs by immunostaining using anti-Dendra2 in Dendra2-NDC80 knock-in line.**

| Sister kinetochore pair # | GSC | SG |
|---|---|---|
| 1 | 1.63255325 | 1.21391 |
| 2 | 2.08886942 | 1.45205 |
| 3 | 1.90246077 | 1.04537 |
| 4 | 1.80469628 | 1.23229 |
| 5 | 1.34713845 | 1.15185 |
| 6 | 1.34905403 | 1.02959 |
| 7 | 1.94231185 | 1.21227 |
| 8 | 1.23244969 | 1.10001 |
| 9 | 1.5934456 | 1.15947 |
| 10 | 1.05108082 | 1.06496 |
| 11 | 1.69266169 | 1.1507 |
| 12 | 3.35622859 | 1.18782 |
| 13 | 2.52883881 | 1.0143 |
| 14 | 2.51928571 | 1.04135 |
| 15 | 3.90118243 | 1.31433 |
| 16 | 1.77010521 | 1.16524 |
| 17 | 1.558123 | 1.00982 |
| 18 | 1.24255173 | 1.19208 |
| 19 | 1.9386881 | 1.1987 |
| 20 | 1.36099849 | 1.17504 |
| 21 | 1.44031101 | 1.21097 |
| 22 | 2.20986391 | 1.21065 |
| 23 | 1.63049841 | 1.28595 |
| 24 | 1.64270188 | 1.15 |
| 25 | 1.50081741 | 1.64763 |
| 26 | 1.97116416 | 1.01702 |
| 27 | 2.83562639 | 1.24521 |
| 28 | 2.10449398 | 1.14968 |
| 29 | 1.39471604 | 1.10928 |
| 30 | 1.3243883 | 1.2998 |
| 31 | 1.21337023 | 1.19974 |
| 32 | 1.23244969 | 1.06658 |
| 33 | 1.5934456 | 1.44734 |
| 34 | 1.25108082 | 1.36097 |
| 35 | 1.29784861 | |
| 36 | 1.81044933 | |
| 37 | 1.94198314 | |
| 38 | 1.36211499 | |
| 39 | 1.52503629 | |

| | |
|---|---|
| **40** | 1.38493551 |
| **41** | 2.072798 |
| **42** | 2.58690102 |
| **43** | 1.37459183 |
| **44** | 1.67051234 |
| **45** | 1.24129449 |
| **46** | 1.34936743 |

**Table S8: Quantification of sister centromeres using anti-CID and sister kinetochores using anti-Dendra2 in Dendra2-NDC80 knock-in line in prometaphase GSCs.**

| **CID and NDC80 at Sister Centromere Pair #** | **CID** | **NDC80** |
|---|---|---|
| **1** | 1.7 | 1.9 |
| **2** | 1.84 | 1.8 |
| **3** | 1.12 | 1.35 |
| **4** | 1.05 | 1.35 |
| **5** | 1.82 | 1.94 |
| **6** | 1.55 | 1.23 |
| **7** | 1.94 | 1.59 |
| **8** | 1.52 | 1.05 |
| **9** | 1.7 | 1.69 |
| **10** | 1.8 | 3.36 |
| **11** | 1.54 | 2.53 |
| **12** | 1.38 | 2.52 |
| **13** | 1.09 | 3.9 |
| **14** | 1.06 | 1.77 |
| **15** | 1.15 | 1.56 |
| **16** | 1.81 | 1.24 |
| **17** | 1.54 | 1.94 |
| **18** | 1.23 | 1.36 |
| **19** | 1.18 | 1.44 |
| **20** | 1.91 | 2.21 |
| **21** | 1.15 | 1.63 |
| **22** | 1.99 | 1.64 |
| **23** | 1.58 | 1.5 |
| **24** | 1.7 | 1.97 |
| **25** | 2.05 | 2.84 |
| **26** | 1.98 | 2.1 |
| **27** | 1.27 | 1.39 |
| **28** | 1.56 | 1.32 |

| | | |
|---|---|---|
| **29** | 1.18 | 1.21 |
| **30** | 1.55 | 1.23 |
| **31** | 1.94 | 1.59 |
| **32** | 1.52 | 1.25 |
| **33** | 1.22 | 1.3 |
| **34** | 1.33 | 1.81 |
| **35** | 1.38 | 1.94 |
| **36** | 1.21 | 1.36 |
| **37** | 1.38 | 1.53 |
| **38** | 1.24 | 1.38 |
| **39** | 1.18 | 2.07 |
| **40** | 1.52 | 2.59 |
| **41** | 1.22 | 1.37 |
| **42** | 1.41 | 1.67 |
| **43** | 1.47 | 1.24 |

**Table S9: Quantification of microtubules% (of the entire microtubules) from mother centrosome *versus* from daughter centrosome using live cell imaging.**

| **Time in min Before metaphase** | **GSC (Mother centrosome)** | **GSC (Daughter centrosome)** | **SG (centrosome 1)** | **SG (centrosome 2)** |
|---|---|---|---|---|
| **00** | 51.96 ± 1.16 | 48.04 ± 1.16 | 50.73 ± 3.74 | 49.27 ± 3.74 |
| **-05** | 43.67 ± 1.81 | 56.33 ± 1.81 | 51.03 ± 4.51 | 48.97 ± 4.51 |
| **-10** | 41.2 ± 2.05 | 58.8 ± 2.05 | 52.55 ± 7.09 | 47.45 ± 7.09 |
| **-15** | 41.35 ± 1.99 | 58.65 ± 1.99 | 53.88 ± 6.85 | 46.12 ± 6.85 |
| **-20** | 41.26 ± 2.18 | 58.74 ± 2.18 | 52.88 ± 6.72 | 47.12 ± 6.72 |
| **-25** | 45.2 ± 2.48 | 54.8 ± 2.48 | 52.81 ± 6.00 | 47.19 ± 6.00 |
| **-30** | 51.21 ± 2.61 | 48.79 ± 2.61 | 52.49 ± 5.82 | 47.51 ± 5.82 |
| **-35** | 53.18 ± 2.62 | 46.82 ± 2.62 | 50.62 ± 5.19 | 49.38 ± 5.19 |
| **-40** | 54.08 ± 2.51 | 45.92 ± 2.51 | 50.73 ± 3.74 | 49.27 ± 3.74 |
| **-45** | 62.16 ± 1.71 | 37.841.71 | | |
| **-50** | 67.94 ± 1.64 | 32.06 ± 1.64 | | |
| **-55** | 73.51 ± 1.56 | 26.491.56 | | |
| **-60** | 77.5 ± 1.23 | 22.51.23 | | |
| **-65** | 76.6 ± 1.12 | 23.41.12 | | |

**Table S10: Quantification of sister centromeres in prometaphase GSCs after *cal1* KD for 5 days (D5) by immunostaining using anti-CID.**

| # Pair of sister centromere | GSC |
|---|---|
| 1 | 1.106504 |
| 2 | 1.103289 |
| 3 | 1.378939 |
| 4 | 1.075039 |
| 5 | 1.319738 |
| 6 | 1.073738 |
| 7 | 1.196998 |
| 8 | 1.760217 |
| 9 | 1.068653 |
| 10 | 1.247975 |
| 11 | 1.001912 |
| 12 | 1.726925 |
| 13 | 1.177936 |
| 14 | 1.691488 |
| 15 | 1.181735 |
| 16 | 1.064402 |
| 17 | 1.114509 |
| 18 | 1.048666 |
| 19 | 1.136748 |
| 20 | 1.279059 |
| 21 | 1.279272 |
| 22 | 1.045063 |
| 23 | 1.122522 |
| 24 | 1.153319 |
| 25 | 1.089298 |
| 26 | 1.230087 |
| 27 | 1.028356 |
| 28 | 1.039924 |
| 29 | 1.075496 |
| 30 | 1.277824 |
| 31 | 1.135423 |
| 32 | 1.244393 |
| 33 | 1.235954 |

**Table S11: Quantification of CID in GSCs in anaphase-early telophase after *cal1* KD for 5 days (D5) by immunostaining using anti-CID.**

| # | GSC |
|---|---|
| **1** | 0.949625921 |
| **2** | 0.917075766 |
| **3** | 1.211244425 |
| **4** | 1.094027837 |
| **5** | 0.857825647 |
| **6** | 1.009131525 |
| **7** | 1.042301184 |
| **8** | 0.9044 |
| **9** | 1.25 |
| **10** | 1.064685908 |
| **11** | 0.916238787 |
| **12** | 1.013377926 |
| **13** | 0.96062992 |
| **14** | 1.253665689 |
| **15** | 1.086193656 |
| **16** | 0.862641927 |

**Table S12: Quantification of number of GSCs after *cal1* KD for 10 days (D10).**

| Testis | Control | *cal1* KD (D10) |
|---|---|---|
| **1** | 10 | 7 |
| **2** | 11 | 6 |
| **3** | 10 | 7 |
| **4** | 7 | 8 |
| **5** | 11 | 7 |
| **6** | 8 | 6 |
| **7** | 7 | 8 |
| **8** | 9 | 7 |
| **9** | 9 | 4 |
| **10** | 9 | 6 |
| **11** | 10 | 8 |
| **12** | 11 | 6 |
| **13** | 10 | 7 |
| **14** | 11 | 5 |
| **15** | 10 | 6 |
| **16** | 11 | 7 |
| **17** | 6 | 5 |
| **18** | 11 | 6 |
| **19** | 7 | 3 |
| **20** | 8 | 6 |
| **21** | 9 | 5 |
| **22** | 10 | 5 |
| **23** | 11 | 6 |
| **24** | 9 | 6 |
| **25** | 9 | 6 |
| **26** | 11 | 6 |
| **27** | 9 | 5 |
| **28** | 11 | 5 |
| **29** | 11 | 5 |
| **30** | 8 | 6 |
| **31** | 7 | 7 |
| **32** | 9 | 5 |
| **33** | 10 | 7 |
| **34** | 11 | 6 |
| **35** |  | 6 |

**Table S13: Quantification of hub area after *cal1* KD for 10 days (D10).**

| Testis | Control ($\mu m^2$) | *cal1* KD ($\mu m^2$, D10) |
|---|---|---|
| **1** | 71.18 | 139.82 |
| **2** | 74.46 | 144.89 |
| **3** | 92.78 | 145.04 |
| **4** | 82.08 | 149.95 |
| **5** | 80.62 | 178.82 |
| **6** | 70.89 | 157.27 |
| **7** | 44.68 | 191.89 |
| **8** | 74.38 | 195.93 |
| **9** | 79.68 | 141.74 |
| **10** | 66.81 | 122.93 |
| **11** | 88.07 | 151.4 |
| **12** | 96.76 | 185.8 |
| **13** | 102.5 | 147 |
| **14** | 70.97 | 155.91 |
| **15** | 82.75 | 102.75 |
| **16** | 87.35 | 91.38 |
| **17** | 99.7 | 123.69 |
| **18** | 92.68 | 150.62 |
| **19** | 60.73 | 194.5 |
| **20** | 98.62 | 141 |
| **21** | 83.58 | 204.53 |
| **22** | 86.78 | 180.75 |
| **23** | 48.33 | 160.49 |

| | | |
|---|---|---|
| **24** | 76.76 | 118.36 |
| **25** | 78.77 | 144.21 |
| **26** | 79.13 | 93.3 |
| **27** | 90.35 | 133.66 |
| **28** | 87.14 | 139.66 |
| **29** | 60.94 | 144.31 |
| **30** | 71.33 | 159.25 |
| **31** | 55.93 | 132.94 |
| **32** | 53.65 | 159.66 |
| **33** | 139.92 | 122.65 |
| **34** | 107.56 | 161.11 |
| **35** | 66.37 | 175.48 |
| **36** | 51.53 | 200.96 |
| **37** | 108.18 | 150.87 |
| **38** | 82.85 | 153.25 |
| **39** | 90.45 | 162.66 |
| **40** | 52.51 | 127.25 |
| **41** | 62.95 | 110.71 |
| **42** | 68.74 | 222.67 |
| **43** | 78.15 | 194.96 |
| **44** | 75.26 | 177.03 |
| **45** | 61.71 | 158.78 |
| **46** | | 158.47 |
| **47** | | 172.79 |
| **48** | | 185.51 |
| **49** | | 108.96 |

| | | |
|---|---|---|
| **50** | | 189.23 |
| **51** | | 217.03 |
| **52** | | 190.67 |
| **53** | | 156.46 |
| **54** | | 153.67 |
| **55** | | 138.83 |

**Table S14: Quantification of microtubules% (of the entire microtubules) from mother centrosome *versus* from daughter centrosome in GSCs after being released from nocodazole treatment using live cell imaging.**

| Time in min (Before metaphase) | GSC (Mother centrosome) | GSC (Daughter centrosome) |
|---|---|---|
| **00** | 52.40 ± 2.24 | 47.59 ± 2.24 |
| **-05** | 49.26 ± 3.09 | 50.74 ± 3.09 |
| **-10** | 51.18 ± 1.62 | 48.82 ± 1.62 |
| **-15** | 51.83 ± 2.44 | 48.17 ± 2.44 |

**Table S15: Quantification of sister centromere in prometaphase immediately after releasing from nocodazole treatment by immunostaining using anti-CID.**

| # Pair of sister centromeres | GSC |
|---|---|
| 1 | 2.103614 |
| 2 | 2.185909 |
| 3 | 1.017633 |
| 4 | 1.075207 |
| 5 | 1.872661 |
| 6 | 1.451219 |
| 7 | 1.267663 |
| 8 | 1.288388 |
| 9 | 1.306424 |
| 10 | 1.05198 |
| 11 | 1.384535 |
| 12 | 1.43337 |
| 13 | 2.017513 |
| 14 | 1.242237 |
| 15 | 1.43811 |
| 16 | 1.736577 |
| 17 | 1.113999 |
| 18 | 2.932079 |
| 19 | 1.948108 |
| 20 | 1.30958 |
| 21 | 2.239273 |
| 22 | 1.087345 |
| 23 | 1.985696 |
| 24 | 1.112867 |
| 25 | 1.373926 |
| 26 | 1.858309 |
| 27 | 2.083878 |
| 28 | 1.223923 |
| 29 | 1.62324 |
| 30 | 1.288315 |
| 31 | 1.298268 |
| 32 | 1.682865 |
| 33 | 1.82598 |
| 34 | 1.293596 |

**Table S16: Quantification of CID signals in anaphase to early telophase GSCs immediately after releasing from nocodazole treatment by immunostaining using anti-CID.**

| # | CID (GSC/GB) |
|---|---|
| 1 | 0.83121 |
| 2 | 0.95958 |
| 3 | 1.01752 |
| 4 | 1.13137 |
| 5 | 1.57944 |
| 6 | 1.07562 |
| 7 | 1.34462 |
| 8 | 0.88964 |
| 9 | 0.98598 |
| 10 | 1.05838 |
| 11 | 1.02209 |
| 12 | 1.00301 |
| 13 | 1.09289 |
| 14 | 1.14886 |
| 15 | 1.14313 |
| 16 | 0.95585 |
| 17 | 1.16803 |
| 18 | 0.87527 |
| 19 | 1.07041 |
| 20 | 1.00645 |
| 21 | 1.05583 |
| 22 | 1.06752 |
| 23 | 1.04396 |
| 24 | 0.94706 |
| 25 | 1.03688 |
| 26 | 1.088333333 |
| 27 | 0.788218794 |
| 28 | 0.748194946 |
| 29 | 1.054867257 |
| 30 | 1.213640099 |
| 31 | 1.400876232 |
| 32 | 1.021494871 |
| 33 | 1.133928571 |
| 34 | 0.964912281 |
| 35 | 0.973214286 |
| 36 | 0.929824561 |
| 37 | 1.396825397 |
| 38 | 0.589928058 |
| 39 | 0.675213675 |
| 40 | 0.848484848 |
| 41 | 1.088607595 |

| | |
|---|---|
| **42** | 0.62 |
| **43** | 0.905660377 |
| **44** | 1.222222222 |
| **45** | 1.240740741 |
| **46** | 0.739130435 |
| **47** | 1.807017544 |
| **48** | 0.706293706 |
| **49** | 1.552631579 |
| **50** | 1.180327869 |
| **51** | 1.044444444 |
| **52** | 0.964912281 |
| **53** | 1.246376812 |
| **54** | 0.961538462 |

**Table S17: Quantification of old and new H3 immediately after releasing from nocodazole in anaphase to early telophase GSCs using fixed cell imaging.**

| # | Old H3 (GSC/GB) | New H3 (GB/GSC) |
|---|---|---|
| **1** | 0.80509 | 0.92097 |
| **2** | 0.84362 | 0.78429 |
| **3** | 0.90605 | 0.86517 |
| **4** | 1.09564 | 0.95767 |
| **5** | 1.00986 | 1.20735 |
| **6** | 0.7556 | 1.01741 |
| **7** | 1.09485 | 1.29103 |
| **8** | 0.94781 | 0.79019 |
| **9** | 0.67165 | 0.69703 |
| **10** | 0.91206 | 1.02308 |
| **11** | 1.01029 | 1.07116 |
| **12** | 1.0613 | 0.93353 |
| **13** | 0.96338 | 1.17816 |
| **14** | 1.03025 | 0.92203 |
| **15** | 1.06317 | 1.20747 |
| **16** | 0.82873 | 0.78745 |
| **17** | 1.0098 | 0.99231 |
| **18** | 1.22729 | 0.89431 |
| **19** | 0.91125 | 0.79001 |
| **20** | 0.89065 | 0.89638 |

**Table S18: Quantification of old H3 immediately after releasing from nocodazole in anaphase GSCs by live cell imaging.**

| # | GSC/GB (WT) | SG1/SG2 (WT) | GSC/GB (immediately after release) | GSC/GB (upon recovery) |
|---|---|---|---|---|
| 1 | 3.00140981 | 1.077560414 | 1.482336224 | 1.77814 |
| 2 | 2.119614267 | 0.942904274 | 0.855046833 | 1.6626 |
| 3 | 2.094502013 | 1.127765327 | 0.944940051 | 1.50504 |
| 4 | 1.710842138 | 1.029377568 | 1.11008235 | 2.38523 |
| 5 | 1.915731947 | 0.988450726 | 0.671087782 | 1.03814 |
| 6 | 1.977206793 | 0.962290083 | 0.901072346 | 1.6753 |
| 7 | 2.119614267 | 0.910348778 | 1.093413811 | 1.63309 |
| 8 | 1.915731947 | 1.251560214 | 0.889588121 | 1.07059 |
| 9 | 1.977206793 | 0.873967438 | 1.057759221 | 1.873 |
| 10 | 1.509965989 | 1.000023552 | 0.782211152 | 1.05179 |
| 11 | 1.559300669 | 1.104569731 | 1.255861906 | 1.40516 |
| 12 | 1.714383102 | 0.948649306 | 0.86744603 | 1.5382 |
| 13 | 2.407387335 | 0.885609811 | 1.289635091 | 1.44014 |
| 14 | 2.008545787 | 1.028150744 | 0.803190272 | 1.25085 |
| 15 | 2.629794826 | 0.987085459 | 1.138100083 | 1.09865 |
| 16 | 2.027497078 | 0.959348448 | 0.883464329 | 2.44607 |
| 17 | 2.094502013 | 0.902657214 | 1.324686941 | 1.41226 |
| 18 | 1.710842138 | 1.033114527 | 1.10417312 | 1.47472 |
| 19 | 1.851770764 | 1.073699216 | 0.9696303 | 1.32155 |
| 20 | 1.606746703 | 1.182152714 | 1.647926736 | 1.03338 |
| 21 | 1.85539857 | 1.171843663 | 1.854211663 | |
| 22 | 2.216178589 | 1.138183797 | 0.998369392 | |
| 23 | 1.743308692 | 1.171843663 | 0.932589688 | |
| 24 | 2.241865072 | 1.164100711 | 1.446077574 | |
| 25 | 2.149067795 | 1.182152714 | 0.85851039 | |
| 26 | 1.531063925 | | 0.875320395 | |
